# PBRM1 BD2 and BD4 associate with RNA to facilitate chromatin association

**DOI:** 10.1101/2022.02.07.479474

**Authors:** Saumya M. De Silva, Surbhi Sood, Alisha Dhiman, Kilsia F. Mercedes, Morkos A. Henen, Beat Vögeli, Emily C. Dykhuizen, Catherine A. Musselman

## Abstract

PBRM1 is a subunit of the PBAF chromatin remodeling complex, which is mutated in 40-50% of clear cell renal cell carcinoma patients. It is thought to largely function as a chromatin binding subunit of the PBAF complex, but the molecular mechanism underlying this activity is not fully known. PBRM1 contains six tandem bromodomains which are known to cooperate in binding of nucleosomes acetylated at histone H3 lysine 14 (H3K14ac). Here we demonstrate that the second and fourth bromodomains from PBRM1 also bind nucleic acids, selectively associating with double stranded RNA elements. Disruption of the RNA binding pocket is found to compromise PBRM1 chromatin binding and inhibit PBRM1-mediated cellular growth effects.

## Introduction

Eukaryotic DNA is packaged into the cell nucleus in the form of chromatin. At its most basic level chromatin is made up of repeats of the basic subunit, the nucleosome(1,2). Each nucleosome is composed of ∼147 base pairs (bp) of DNA, wrapped around an octamer of histone proteins containing two copies each of H2A, H2B, H3, and H4(3). Spatial and temporal modulation of chromatin structure is critical in all DNA-templated processes. This is facilitated by several mechanisms including ATP-dependent remodeling of the nucleosome by multi-subunit protein complexes known as chromatin remodelers(4,5). The action of the remodelers is tightly regulated by a variety of factors including complex composition and post-translational modification (PTM) of the histone proteins(6), and can also be modulated by association with non-coding RNAs (ncRNA)(7,8). Together, these factors stabilize complexes at specific regions of chromatin as well as modulate their activity.

There are several families of chromatin remodeling complexes, including SWI/ SNF, which was first identified in yeast(9). In humans, the SWI/SNF family of remodelers exist primarily as cBAF (hSWI/SNF-A) (10–12), PBAF (hSWI/SNF-B)(13,14), and GBAF (ncBAF)(15,16). PBAF is defined by the exclusive incorporation of ARID2, BRD7, BAF45A and PBRM1 (also known as BAF180 or PB1)(17,18). SWI/SNF subunits are mutated in ∼20% of human tumors(19), with the PBRM1 subunit mutated primarily in 40-50% of clear cell renal cell carcinomas (ccRCC)(20,21). PBRM1 is a bona fide tumor suppressor in ccRCC in the context of VHL deletion(22–24); however, in other cancer contexts, it is oncogenic and a proposed therapeutic target(25–27). Though its best characterized function is in DNA damage repair(28), its role in oncogenesis is also associated with transcriptional activity(29,30), at least in part through the activation of stress-response genes(31,32).

PBRM1 contains several chromatin-binding domains and is thought to function largely as a chromatin targeting subunit of PBAF. This includes six bromodomains (BDs), two bromo-adjacent homology (BAH) domains, and a high mobility group (HMG) domain (Figure 1a)(14,33). The HMG domain of PBRM1 has been shown to interact with nucleosomal DNA(10,34,35), whereas the BAH domains are thought to be protein-protein interaction domains(36), with one recent study implicating them as readers for methylated lysine 40 of α-tubulin on spindle microtubules(37). Bromodomains (BDs) are ∼100aa four-helix bundles that associate with acetylated histones(38– 40). The BDs of PBRM1 are structurally well conserved despite substantial differences in sequence (Supplementary Figure 1). Studies on the six BDs reveal that they cooperate in binding to nucleosomes containing histone H3 acetylated at lysine 14 (H3K14ac)(41–43). From several different assays BD2 and BD4 appear to be most important for robust H3K14ac binding with BD1, BD5, and BD6 enhancing activity, while BD3 has been proposed to attenuate it. The acetyl-lysine binding activity of the BDs, particularly BD2 and BD4, are critical for cell proliferation and PBRM1-mediated gene expression(42–44).

**Figure 1.**
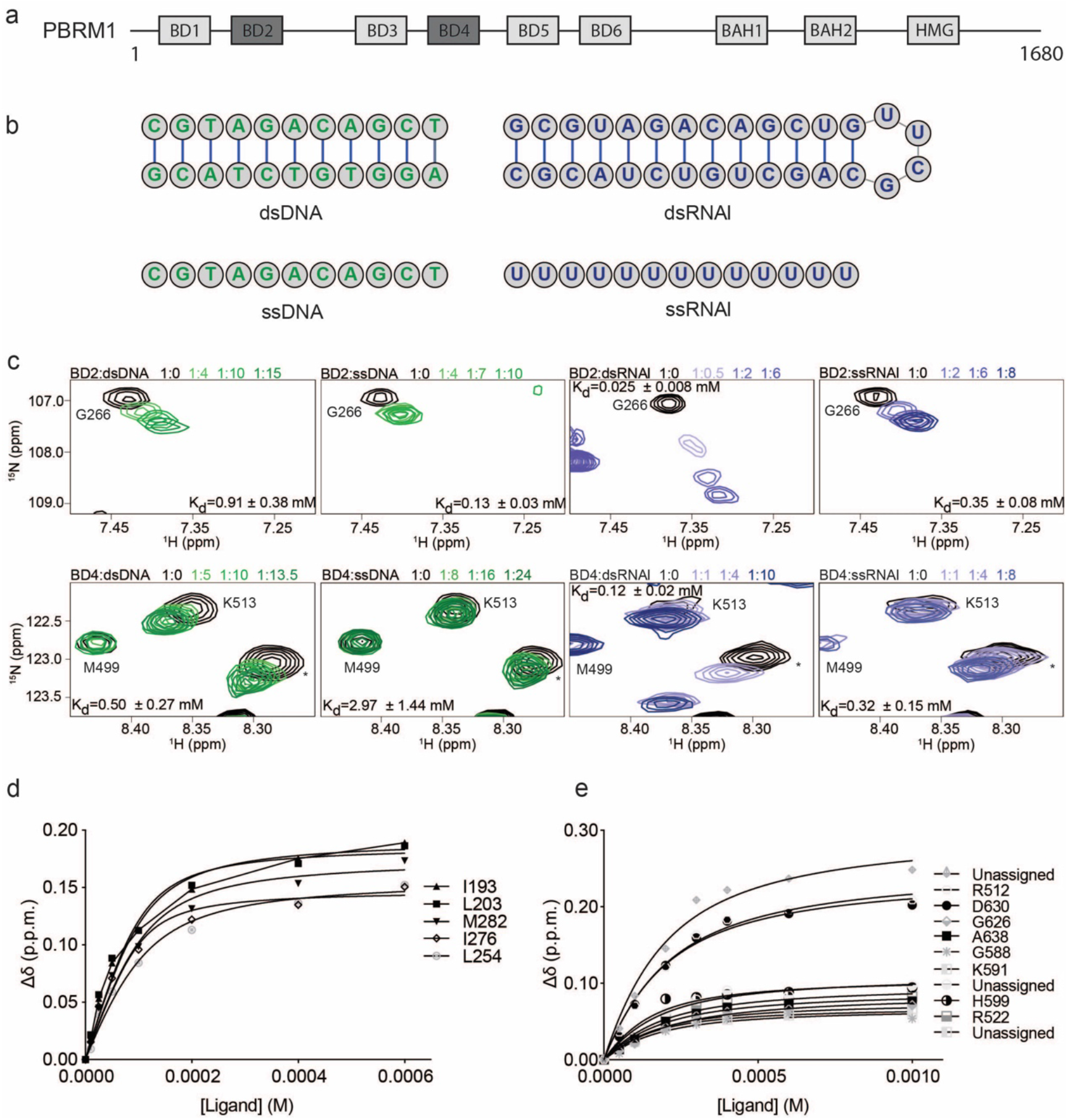
PBRM1 BD2 and BD4 selectively bind to RNA. **(a)** The domain architecture of the PBRM1 subunit of PBAF complex. BD2 and BD4 are colored dark grey. **(b)** Structures of double stranded (top) and single stranded (bottom) DNA (green) and RNA (purple) used for NMR titrations shown in c. **(c)** Overlay of ^1^H-^15^N HSQC spectra of ^15^N-BD2 (top row) or ^15^N-BD4 (bottom row) upon titration with dsDNA, ssDNA, dsRNAI and ssRNAI. The spectra are color coded according to protein:nucleic acid molar ratio as shown in the legend. For clarity, only 4 titration points are displayed. For BD2 dsDNA titration was collected at 1:0, 1:1, 1:2, 1:4, 1:6, 1:10, 1:12, 1:15, ssDNA was collected at 1:0, 1:0.5, 1:1, 1:2, 1:4, 1:7, 1:10, dsRNAI was collected at 1:0, 1:0.1, 1:0.25, 1:0.5, 1:1, 1:2, 1:4, 1:6 and ssRNAI was collected at 1:0, 1:1, 1:2, 1:4, 1:6, 1:8. For BD4 sDNA titration was collected at 1:0, 1:0.5, 1:1, 1:2, 1:2.5, 1:5, 1:7.5, 1:10, 1:13.5, ssDNA was collected at 1:0, 1:0.5, 1:1, 1:4, 1:8, 1:12, 1:16, 1:24, dsRNAI was collected at 1:0, 1:0.5, 1:1, 1:2, 1:3, 1:4, 1:6, 1:10 and ssRNAI was collected at 1:0, 1:1, 1:2, 1:4, 1:6, 1:8. **(d**,**e)** Binding curves calculated from the normalized chemical shifts for BD2 (d) or BD4 (e) upon binding to dsRNAI. Titration points are fit to a single-site binding model under ligand-depleted conditions.

Recently, a subset of BDs has been discovered to associate with DNA or RNA in addition to binding acetylated histone tails(45–50). The BRG1 and hBRM BDs as well as BDs in BRD2, BRD3, BRD4, and BRDT interact with free and nucleosomal DNA. In addition, BRD3 and BRD4 BDs associate with non-coding RNA (ncRNA)(49,50). While the DNA and acetylated histone interactions have been found to be largely independent, there is evidence that RNA and acetylated histone interactions are cooperative(47), though the mechanism for this is unknown.

Morrison et. al predicted that ∼30% of known BDs will bind to nucleic acids, and this includes BD2, BD3, BD4, and BD5 of PBRM1(46). Here we demonstrate that PBRM1 BD2 and BD4 selectively associate with double-stranded RNA elements, this binding enhances the association with H3K14ac peptides, and the nucleic acid binding activity of these domains is important for PBRM1-mediated chromatin association and cell proliferative effects.

## Methods

### Cloning and Expression of PBRM1 Bromodomains and Bromodomain Mutants

Codon optimized plasmids encoding the individual His-tagged PBRM1 bromodomain constructs were received from Nicola Burgess-Brown (Addgene plasmid numbers 38999, 39013, 39027, 39028, 39030, 39103). Individual BDs were as follows: BD1 (residues 43-156), BD2 (residues 178-291), BD3 (residues 388-494), BD4 (residues 496-637), BD5 (residues 645-766), BD6 (residues 774-917). Bromodomain mutants were generated using the Q5 site-directed mutagenesis kit (New England Biolabs) and each mutation was confirmed through DNA sequencing. All constructs were expressed in *E. coli* BL21 DE3 cells (New England Biolabs, Ipswich, MA). Cells were grown in LB media (VWR) at 37 °C in the presence of 50 μg/L kanamycin (VWR). When the cultures reached an OD_600_ ∼1 – 1.2, cell were harvested at 4,000 rpm for 15 minutes at 22 °C. Cell pellets were resuspended in M9 minimal media (at 1:4 ratio of M9:LB) containing 1g/L ^15^N ammonium chloride and 5g/L D-glucose or 4g/L ^13^C-D-glucose, as well as vitamins (Centrum, 5 mL of 1 tablet resuspended in 50 mL water). After resting for an hour, cells were induced with 1 mM IPTG and allowed to grow at 18 °C for 18 hours. Cells were harvested by centrifugation at 6,000 rpm for 20-30 minutes at 4 °C. Cell pellets were flash frozen in liquid N_2_ and stored at −80 °C.

### Protein Purification

Cell pellets were resuspended in lysis buffer containing 50 mM potassium phosphate at pH 7.0, 500 mM KCl, 2 mM DTT, 0.5% Triton X-100, 0.5 mg ml^-1^ lysozyme, DNAse I (RNase free – Thermo Fisher), and an EDTA-free protease inhibitor cocktail tablet (Roche mini tablets). Cells were lysed via sonication and the cell lysate was cleared by centrifugation at 15,000 rpm at 4 °C for one hour. The soluble fraction was loaded onto a column packed with Ni(II)-nitriloriacetic acid agarose (Qiagen) pre-equilibrated in 100 mL of binding buffer (50 mM potassium phosphate at pH 7.0, 50 mM KCl, 2 mM DTT and 5 mM imidazole), then eluted with 200 mM-500 mM imidazole and dialyzed against dialysis buffer to remove imidazole. The His tag was cleaved using TEV protease at 18 °C for 4-6 hours. Cleaved protein was further purified by fast protein liquid chromatography (FPLC) using a Source-S cation-exchange column (GE Healthcare) (low salt buffer with 50 mM potassium phosphate at pH 7.0, 50 mM KCl, 2 mM DTT, 2 mM EDTA and high salt buffer with 50 mM potassium phosphate at pH 7.0, 1M KCl, 2 mM DTT, 2 mM EDTA). This was followed by purification over a Superdex 75 size-exclusion column (GE Healthcare). The final buffer was 50 mM potassium phosphate at pH 7.0, 50 mM KCl, 2 mM DTT, 2 mM EDTA. For PBRM1 BD2 and BD4, there was incomplete cleaving of the His tag. Therefore, after cleavage the protein was run over a Ni(II)-nitriloriacetic acid agarose column again before the cation exchange step to remove the un-cleaved species.

### Electrophoretic Mobility Shift Assays (EMSAs)

601 DNA was purified as previously described(51). Samples for EMSAs were prepared by mixing 1.5 pmol of Widom 601 DNA with varying amounts of individual bromodomains to reach final molar ratios of DNA:protein of 1:0, 1:1, 1:4, 1:6, 1:10, 1:15, 1:20 and 1:30 in a final buffer containing 50 mM potassium phosphate at pH 7.0, 10 mM KCl, 2 mM DTT, 2 mM EDTA and 5% v/v sucrose for gel loading purposes. While the samples were equilibrating on ice for an hour 5% 75:1 acrylamide:bisacrylamide native gel was pre-run in 0.2X TBE buffer on ice at 4 °C for 30 minutes at 120 V. The samples were run on ice at 4 °C for 50 minutes at 125 V. Gels were stained with ethidium bromide or SYBR green (Fisher) and visualized using an ImageQuant LAS 4000 imager/BIO-RAD Gel Doc imager.

### Preparation of oligonucleotides

DNA and RNA oligonucleotides for NMR experiments were obtained from Integrated DNA Technologies (IDT, Inc.). The following sequences were used: dsDNA (5’ CGTAGACAGCT 3’ and 5’ AGCTGTCTACG 3’), ssDNA (5’ CGTAGACAGCT 3’), dsRNAI (5’ GCGUAGACAGCUGUUCGCAGCUGUCUAUGC 3’), dsRNAII (5’ GGCAUCGUGCUUC GGCACGAUGCC 3’), dsRNAIII (5’ GGCAUCGUGCAUCAGAAUGGCACGAUGCC 3’), dsRNA IV (5’ GGCAUCGUGC 3’ and 5’ GCACGAUGCC 3’), ssRNAI (5’ UUUUUUUUUUUUU 3’), ssRNAII (5’ ACACACACACACA 3’). To anneal the DNA, single stranded oligos were mixed at 1:1 molar ratio, heated to 95 °C for 5 minutes and allowed to cool overnight to room temperature. To anneal or refold RNA, oligonucleotides were dissolved in 50 mM potassium phosphate at pH 7.0, 50 mM KCl, 2 mM DTT, 2 mM EDTA, heated to 95 °C for 5 minutes and allowed to cool on ice for one hour. All of these were further purified by FPLC using a Superdex 75 column (GE Healthcare) in a buffer containing 50 mM potassium phosphate at pH 7.0, 50 mM KCl, 2 mM DTT, 2 mM EDTA and concentrated. Single stranded nucleic acids were dissolved in 50 mM potassium phosphate at pH 7.0, 50 mM KCl, 2 mM DTT, 2 mM EDTA to get the final desired concentration. At the concentrations used for NMR titrations, the dsDNAs and dsRNAs are predicted to be at least 98% in duplex according to IDT.

### Preparation of Histone Tail Peptides

Histone peptide corresponding to H3K14Ac (9-19) was obtained from Bon Opus. All histone peptides were dissolved in H_2_O at a concentration of 20 mM determined by manufacturer determined weight, and the pH adjusted to ∼7.0.

### NMR spectroscopy and data analysis

NMR experiments were carried out on either an 800 MHz Bruker Avance II, Bruker Avance Neo 600 MHz, Bruker Avance II 500 MHz, Varian INOVA 600 MHz, or Varian 900 MHz all equipped with cryoprobes. To obtain backbone assignments for BD2 and BD4, HNCACB, CBCA(CO)NH, HNCO and HN(CO)CA(52) (BD2 only) spectra were collected on ^15^N-^13^C-labelled sample at 1mM on a Bruker 500 MHz (for BD4) and Bruker Avance Neo 600 MHz (for BD2) at 25 °C. NMR data were processed using NMRPipe(53) and analyzed in CcpNmr(54). For the experiments carried out using nonuniform sampling, spectra were reconstituted with SMILES(55).

Titrations of wild-type and mutant BDs with oligonucleotides or acetylated peptides were carried out by collecting ^1^H,^15^N heteronuclear single quantum coherence (HSQC) spectra on ^15^N-labelled BDs at 0.1 mM in 93% H_2_O/7% D_2_O at 25 °C upon addition of increasing concentrations of substrate. NMR data were processed in NMRPipe(53) and analyzed in CcpNmr(54). The normalized chemical shift difference (Δδ) at every titration point was calculated as

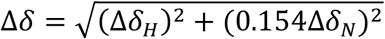

where Δδ_H_ and Δδ_N_ are the change in the ^1^H and ^15^N chemical shift respectively between the apo state and each titration point. Dissociation constants (K_d_s) were calculated by fitting the data to a single-site binding model accounting for ligand depletion in GraphPad Prism using the equation(56)

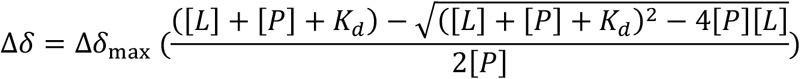

where [L] is the concentration of ligand, [P] is the concentration of the bromodomain and Δδ_max_ is the chemical shift change at saturation. Reported K_d_ values were calculated as an average of individual values for residues with significant Δδ for each individual titration. Residues were determined to be significantly perturbed, if Δδ was greater than average Δδ plus 2 standard deviations of all residues (excluding unassigned residues) after trimming the 10% of residues with the largest Δδ.

### Cell culture conditions

Caki2 cells (American Type Culture Collection, Manassas, VA) were grown in McCoy’s 5A medium (Corning Mediatech, Inc., Manassas, VA) supplemented with 10% fetal bovine serum (Corning Mediatech, Inc., Woodland, CA), 100 units/ml penicillin and 100 μg/ml streptomycin (Corning Mediatech, Inc.), 1% MEM nonessential amino acids (Corning Mediatech, Inc.), and 2 mM L-alanyl-L-glutamine (Corning Glutagro; Corning Mediatech, Inc.) at 37 °C in a humidified atmosphere in a 5% CO_2_ incubator.

### Generation of cell lines expressing full length PBRM1 mutants

A section of full length *PBRM1* (nucleotides 434-2130) with triple mutations (S275A,K277A,Y281A-SKY, K511A,R512A,K513A-KRK) were purchased from Biomatik and cloned into BstBI-digested PBRM1-TetO-FUW using the ligation-free In-Fusion HD cloning kit (Takara). The construct was packaged into lentivirus using HEK293T cells and delivered into target cells together with pLenti CMV rtTA3 Hygro (w785-1) (a gift from Eric Campeau Addgene plasmid no. 26730) for tetracycline inducible expression. The cells were selected with puromycin (2 μg/mL) and hygromycin B (200 μg/mL). To induce PBRM1 expression, Caki2 cells were treated with doxycycline (2 μg/mL) for at least 3 days.

### Nuclear Lysates

Caki2 cells were tryspinized and washed in ice-cold phosphate buffered saline (PBS at pH 7.2). The pellet was resuspended in buffer A (20 mM HEPES at pH 7.9, 25 mM KCl, 10% glycerol, 0.1% Nonidet P-40 with protease inhibitors) at a concentration of 20 million cells/ml. The cells were kept on ice for 5 minutes, and nuclei were isolated by centrifugation at 600 × *g* (Eppendorf Centrifuge 5810 R, Hamburg, Germany) for 10 minutes. The nuclei pellet was resuspended in lysis buffer (50 mM Tris at pH 7.4, 150 mM NaCl, 1% Nonidet P-40, 1 mM EDTA, and protease inhibitors) at a concentration of 50 million nuclei/mL and rotated at 4 °C for 30 min. The lysate was cleared by centrifugation at 20,000 x g for 10 min.

### Immunoblot analysis

Protein samples were mixed with 4x lithium dodecyl sulfate sample buffer containing 10% 2-merchaptoethanol. The protein lysates were denatured for 5 minutes at 95 °C, separated on a 4– 12% SDS-polyacrylamide gel, and transferred to a PVDF membrane (Immobilon FL, EMD Millipore, Billerica, MA). The membrane was blocked with 5% bovine serum albumin (VWR, Batavia, IL) in PBS containing 0.1% Tween-20 for one hour at room temperature and then incubated in primary antibodies overnight at 4 °C. The primary antibodies were detected by incubating the membranes in goat anti-rabbit or goat anti-mouse secondary antibodies (LI-COR Biotechnology, Lincoln, NE) conjugated to IRDye 800CW or IRDye 680CW, respectively, for one hour at room temperature, and the signals were visualized using Odyssey Clx imager (LI-COR Biotechnology). Antibodies used for immunoblot: PBRM1 (Bethyl, A301-590A, 1:1000), ARID1A (Santa Cruz 1:1000) V5 (CST, D3H8Q, 1:1000), hnRNP A1 (Santa Cruz, 4B10 1:1000).

### Growth curve analysis

For growth curve analysis, 2,000 (cell number) control or single/ triple mutant cells were plated in 96-well plates. After 7 days, culture medium was aspirated and percent viability was assessed using CellTiter-Glo® reagent (Promega, Madison, WI). The luminescence was measured using Promega™ Glomax® luminometer.

### Cell Proliferation competition assay

Caki2 parental cells were first transduced with lentiviral particles for the dual reporter construct pFU-Luc2-eGFP (L2G)(57) (a kind gift from Huiping Lui) and the GFP expressing cells were selected using FACS. These GFP positive cells were transduced with lentiviral particles for pLenti CMV rtTA3 Hygro (w785-1) (a gift from Eric Campeau Addgene plasmid number 26730) for tetracycline inducible expression and selected using hygromycin (200μg/ml). Once selected, these cells were transduced with lentiviral particles for TetO-Fuw empty vector (Addgene plasmid number 85747)(30). This cell line is referred to as Caki2 GFP-Fuw in the later sections.

Caki2 parental cells were transduced with rtTA and either TetO-Fuw empty vector, TetO-Fuw-PB1 WT (Addgene plasmid # 85746)(30), TetO-Fuw-PB1-SYK, or TetO-Fuw-PB-KRK. All Caki2 cells were induced for protein expression using 2 μg/ml of doxycycline for 72 hours, following which they were seeded in 1:1 ratio in a 6-well plate and cultured in the presence of 2 μg/ml of doxycycline throughout the duration of the experiment. At 24 hour post-seeding, each well was trypsinized and 1/4 of the harvested cells were re-seeded in a 6-well for the next timepoint while 3/4 of the harvested cells were analyzed by flow cytometry to determine GFP+ and GFP-populations. The 24 hour GFP-/GFP+ ratio was used as a baseline for all the subsequent timepoints. The co-culture wells were harvested every 72-96 hours depending on the confluency, such that the confluency never crossed 70%. Cell populations were analyzed using the Guava EasyCyte benchtop flow cytometer using monoculture cells as the controls and data analysis was done using FlowJo and GraphPad Prism.

### Serial salt extraction assay

Serial salt extraction assay was performed as published with minor modifications(58). Briefly, 5 million Caki2 cells were harvested by trypsinization and washed once with ice-cold PBS. The cells were lysed in modified buffer A (60 mM Tris, 60 mM KCl, 1 mM EDTA, 0.3 M sucrose, 0.5% Nonidet P-40, 1 mM DTT) with protease inhibitor and deacetylases inhibitor, and nuclei were pelleted. The nuclei were then incubated in 200 μl of extraction buffer 0 (50 mM HEPES at pH 7.8, 2% NP-40, 0.5% sodium deoxycholate, 1mM DTT, protease inhibitors and deacetylase inhibitor) for 10 minutes and centrifuged at 7,000 × *g* for 5 minutes, and supernatant was collected as “0 mM fraction”. The pellet was then resuspended in 200 μl of extraction buffer 100 (50 mM HEPES at pH 7.8, 2% NP-40, 0.5% sodium deoxycholate,1 mM DTT, protease inhibitors, 100 mM NaCl) and processed in the same manner to yield “100 mM fraction.” Serial extraction was implemented with extraction buffers containing 200, 300, 400, and 500 mM NaCl. 20 μl aliquots from each fraction were mixed with 4× lithium dodecyl sulfate loading buffer and run for Western blotting.

### Cross Linking Immunoprecipitation (CLIP)

Caki2 cells were washed in ice-cold phosphate buffered saline (PBS at pH 7.2). The cells were coated with PBS and irradiated at 200 mJ/cm^2^ using SpectroLinker XL_1000 UV crosslinker. The pellet was resuspended in buffer A (20 mM HEPES at pH 7.9, 25 mM KCl, 10% glycerol, 0.1% Nonidet P-40 with PMSF, aprotinin, leupeptin, and pepstatin, DTT, SAHA) at a concentration of 20 million cells/ml. The cells were kept on ice for 5 minutes, and nuclei were isolated by centrifugation at 600 × *g* (Eppendorf Centrifuge 5810 R, Hamburg, Germany) for 10 minutes. The nuclei pellet was resuspended in RIPA buffer (50 mM Tris at pH 7.4, 150 mM NaCl, 1% Nonidet P-40, 0.1% SDS, 0.5% sodium deoxycholate supplemented with PMSF, aprotinin, leupeptin, and pepstatin, DTT, SAHA) and incubated on ice for 5 minutes. 1:50 dilution of RNase cocktail (Ambion, Inc., Foster City, CA; 500 U/mL RNase A+20,000 U/mL RNase T1) and 4U of Turbo DNase (Invitrogen, 2 U/μL) were added to extracts and incubated at 37 °C for 3 minutes. The extracts were cleared by centrifugation (Centrifuge 5424 R; Eppendorf, Hamburg, Germany) at 21,000 × *g* for 30 minutes. One microgram specific antibody was used per 200 μL lysate for immunoprecipitation V5 (CST, D3H8Q, 1:200 IP), or hnRNP A1 (Santa Cruz, 4B10 1:200 IP).

After 1 hour incubation, immunocomplexes were captured using protein A magnetic beads following a 2 hour incubation. The beads were washed twice in high salt buffer (50 mM Tris at pH 7.9, 1 M NaCl, 1 mM EDTA, 1M Urea, 1% Nonidet P-40, 0.1% SDS, 0.5% sodium deoxycholate, with PMSF, aprotinin, leupeptin, and pepstatin, DTT, SAHA, ribo-out) followed by two washes in low salt buffer (20 mM Tris at pH 7.9, 0.2% Tween-20, 10 mM MgCl_2_ with PMSF, aprotinin, leupeptin, and pepstatin, DTT, SAHA, ribo-out). The beads were resuspended in 500 μL of low salt buffer and divided 1:3 for western blot and PNK labeling. The western blot protein samples were eluted in 1× lithium dodecyl sulfate loading dye (Thermo Scientific) by boiling at 95 °C for 5 minutes. For PNK labeling, PNK enzyme (Thermo Scientific, VA) and γ ATP (PerkinElmer) were added to the beads and incubated at 37 °C for 30 minutes. The hot reaction mix was removed, the beads were boiled in 1X loading dye, the proteins were separated on a 4– 12% SDS-polyacrylamide gel and labeled proteins were imaged using a phosphorimager (Typhoon ® FLA9500).

## Results

### PBRM1 BD2 and BD4 selectively associate with RNA

To initially test the nucleic acid binding activity of PBRM1 BDs, electromobility shift assays (EMSAs) were performed for each BD individually with the 147 base-pair (bp) Widom 601 DNA. Consistent with our previous hypothesis(46), the addition of increasing concentrations of BD2, BD3, BD4, and BD5 resulted in a shift in the migration of DNA, indicative of binding (Supplementary Figure 2). In contrast, no shift was observed for BD1 and BD6 at any of the concentrations tested, consistent with no to very weak DNA binding of these BDs (Supplementary Figure 2).

To further characterize these interactions, we utilized nuclear magnetic resonance (NMR) spectroscopy. Initial ^1^H,^15^N heteronuclear single quantum coherence (^1^H,^15^N-HSQC) spectra of each BD revealed well-dispersed resonances in both ^1^H and ^15^N dimensions, indicating that all domains are well-folded (Supplementary Figure 3). However, BD3 and BD5 were unstable over time (data not shown), and as such, we focused all further studies on BD2 and BD4.

To assess the nucleic acid selectivity of BD2 and BD4, we tested binding to four nucleic acid substrates: 11 base pair double-stranded DNA (dsDNA), 13 nucleotide (nt) single-stranded DNA (ssDNA), 30nt stem-loop RNA (dsRNAI), and 13nt poly-U ssRNA (ssRNAI) (see Figure 1b). ^1^H-^15^N HSQC spectra were collected on ^15^N-labeled BD2 or BD4 upon titration of each oligonucleotide substrate (Figure 1c, Supplementary Figures 4,5). Chemical shift perturbations (CSPs) were observed in BD2 and BD4 resonances upon increasing concentrations of all substrates tested, consistent with binding; however, differences in the patterns of CSPs suggest clear differences in the strength of interaction. Dissociation constants were calculated by plotting the normalized CSPs as a function of substrate concentration and fitting to a single-site binding model accounting for ligand depletion (see methods). This revealed that BD2 has the highest affinity for the stem-loop dsRNAI (K_d_ = 0.025 mM ± 0.008 mM, Figure 1d) compared to substantially weaker affinity for dsDNA (K_d_ = 0.91 mM ± 0.38 mM), ssRNAI (K_d_ = 0.35 mM ± 0.08 mM) and ssDNA (K_d_ = 0.13 mM ± 0.03 mM). BD4 also demonstrated the highest affinity for the stem-loop dsRNAI, albeit with weaker overall affinity as compared to BD2 (K_d_ = 0.12 mM ± 0.02 mM, Figure 1e). In comparison, dsDNA bound with K_d_ = 0.50 mM ± 0.27 mM, and ssDNA and ssRNAI bound with K_d_ = 2.97 mM ± 0.14 mM and K_d_ = 0.32 mM ± 0.14 mM, respectively.

To better understand the selectivity for the stem-loop RNA, we investigated additional RNA substrates, including two additional stem-loop structures and a dsRNA lacking any loop (Figure 2c). CSPs were observed in both BD2 and BD4 upon titration of all dsRNA substrates (Figure 2 a,b, Supplementary Figures 4,5). Notably, for the majority of residues, the CSPs follow the same trajectory for all stem-loops, as well as the dsRNA lacking a loop. The exception is M282 of BD2, which has slightly different trajectories for dsDNA, stem-loop RNAs, and dsRNA lacking the loop. In addition, the calculated K_d_ values were similar for all dsRNA substrates for both BDs, but overall weaker for BD4 as compared to BD2 (Figure 2d). Together, this suggests that the association of BD2 and BD4 with RNA are not dependent on the loop of the BD, and rather that both BDs are selectively interacting with double-stranded RNA elements.

**Figure 2.**
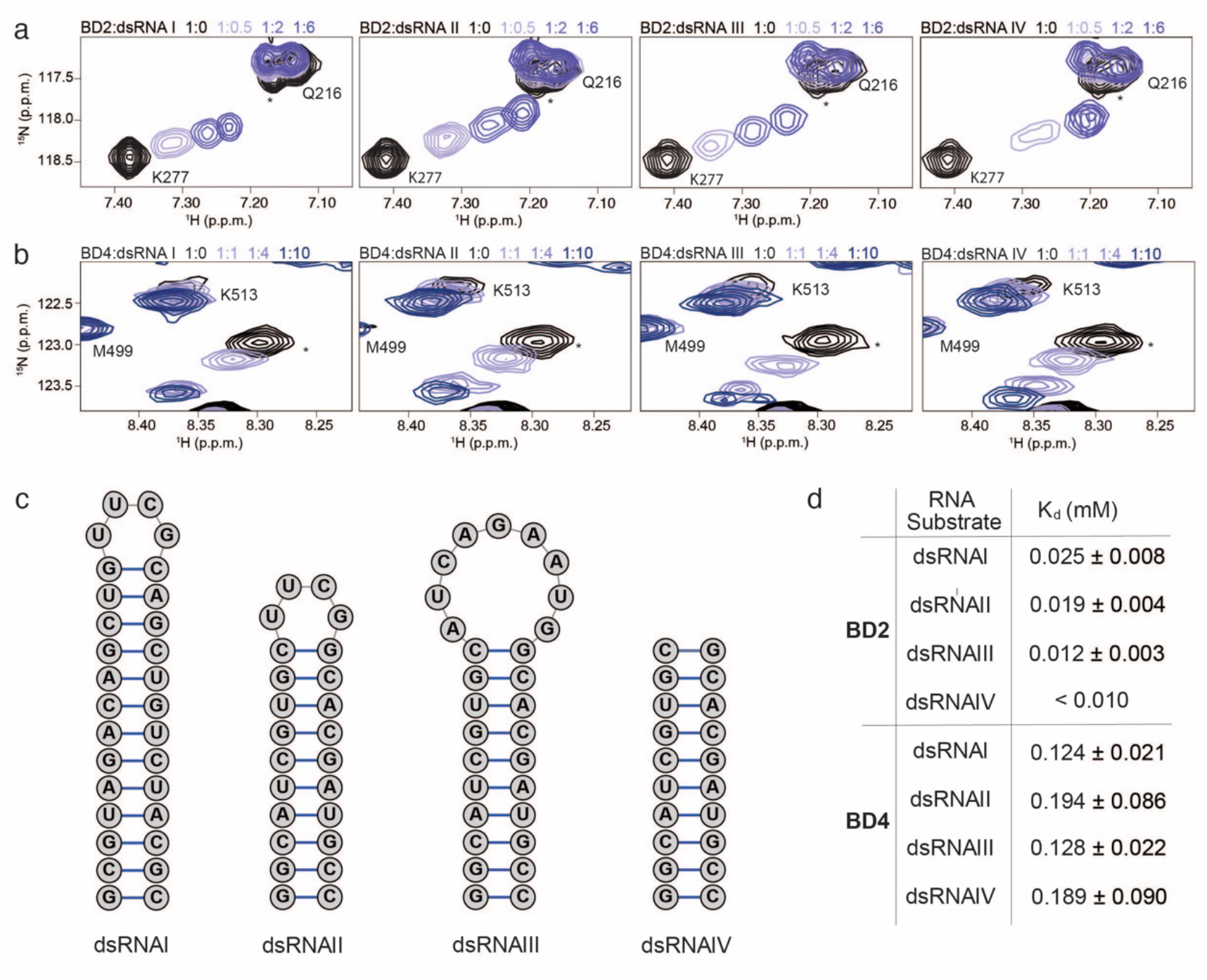
BD2 and BD4 preferentially associate with double stranded RNA elements. **(a**,**b)** Overlay of ^1^H-^15^N HSQC spectra of ^15^N-BD2 (a) or ^15^N-BD4 (b) upon titration with different dsRNA substrates (see c). The spectra are color coded according to protein:RNA molar ratio as shown in the legend. For clarity, only 4 points are displayed. For BD2, all dsRNA titrations were collected at 1:0, 1:0.1, 1:0.25, 1:0.5, 1:1, 1:2, 1:4, 1:6. For BD4, all titrations were collected at 1:0, 1:0.5, 1:1, 1:2, 1:3, 1:4, 1:6, 1:10. **(c)** Schematics of dsRNA substrates used for NMR titrations in the study (d) Dissociation constants (K_d_) determined from NMR titrations with ^15^N-PBRM1 BD2 and BD4 with double stranded RNA substrates. K_d_ values are fit to a single-site binding model under ligand-depleted conditions using 7-9 titration points. Shown are the averages over individual residues with significant CSPs and associated standard deviation. For dsRNAIV association with BD2 the calculated K_d_ was less than 1/10 the protein concentration and thus stated as the upper limit.

### BD2 and BD4 bind RNA through distinct binding pockets

RNA binding pockets on BD2 and BD4 were identified by plotting the normalized CSPs as a function of BD residue. For BD2, association with dsRNAI leads to the largest CSPs in residues in the BC loop and αC helix with smaller perturbations observed in the αB helix and ZA loop (Figure 3a). Notably, all three stem-loop RNAs and the dsRNA with no loop demonstrated nearly identical binding pockets on BD2, suggesting an identical mechanism of binding to these substrates and consistent with the loop playing no role in the interaction (Supplementary Figure 6). Mapping the CSPs onto a previously solved structure of BD2 reveals that these residues cluster along one face of the alpha-helical bundle formed by αB and αC, that coincides with a highly basic surface patch (Figure 3c and Supplementary Figure 7a).

**Figure 3.**
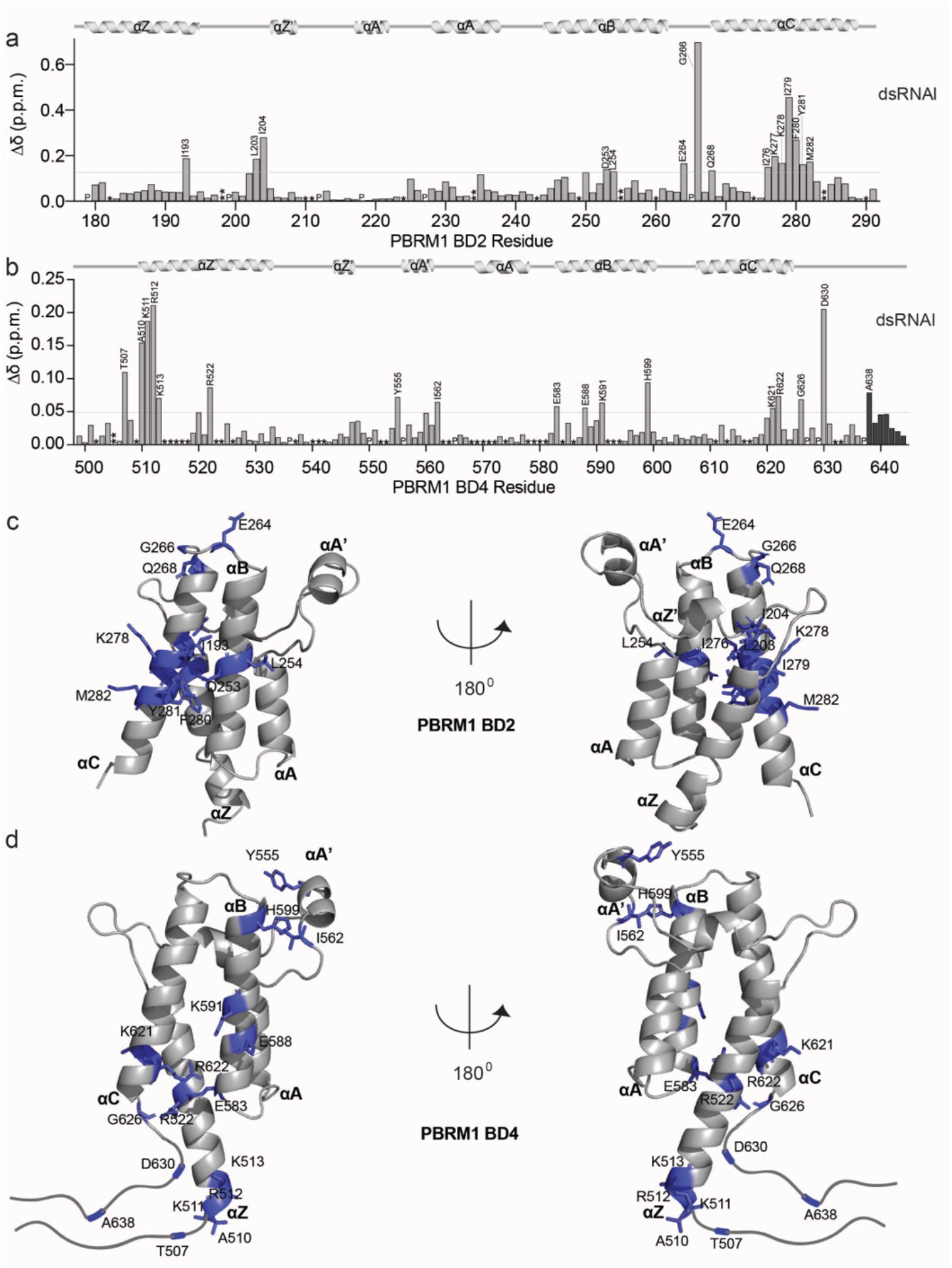
BD2 and BD4 bind RNA through distinct binding pockets. **(a**,**b)** Normalized chemical shift changes between apo and RNA-bound (Δδ) plotted as a function of residue for BD2 (a, 1:6 of BD2:dsRNAI) or BD4 (b, 1:10 of BD4:dsRNAI). Prolines are marked as (P), residues that are unassigned in the apo are marked as (*), residues for which the bound states were not trackable due to overlap are marked as (**), residues that broadened beyond detection are marked as (***). The secondary structure of the BDs is denoted above the plots, and residues that were perturbed greater than the average plus two standard deviations (grey line) after trimming off the top 10% are labelled. **(c**,**d)** Residues significantly perturbed upon addition of dsRNAI are shown as purple sticks and labeled on the previously solved structure of BD2 (c, PDB ID 3HMF) or BD4 (d, PDB ID 3TLP) with the secondary structure elements labelled.

A similar analysis for BD4 reveals a distinct binding pocket with the most perturbed residues in a region N-terminal to the αZ helix as well as in αZ, and similarly in the residues C-terminal to the αC helix and in αC. Smaller perturbations are also observed in the αB helix and ZA loop (Figure 3b). Mapping these onto a previously solved crystal structure of BD4 reveals a much more extended binding pocket as compared to BD2, spanning from the bottom of the alpha-helical bundle to the ZA loop, but also coinciding with an elongated basic patch (Figure 3c and Supplementary Figure 7b). The residues at the N- and C-termini were not included in the construct used for the crystal structure. However, analysis of the HN, N, Cα, Cβ, and CO chemical shifts using TALOS+(59) (Supplementary Figure 7c) or a calculated SSP score (60) (Supplementary Figure 7d) indicate that these residues are largely disordered though with some helical propensity in both N and C termini. As observed for BD2, all three stem-loops and the dsRNA without a loop interact with nearly identical binding pockets on BD4 (Supplementary Figure 8).

For BD2, ssRNA, ssDNA, and dsDNA substrates led to CSPs in nearly identical residues as compared to dsRNA but led to substantially lower magnitudes of CSPs consistent with a less stable complex (Supplementary Figure 6,8). A similar effect is seen for BD4 with ssRNA and dsDNA. Notably, BD4 has some additional residues that are perturbed upon addition of ssDNA but overall the magnitude of CSPs are still quite small as compared to RNA (Supplementary Figure 8).

### RNA association enhances histone tail binding

Both BD2 and BD4 have previously been shown to preferentially associate with H3K14ac peptides and are the most critical BDs for PBRM1 association with H3K14ac nucleosomes(43). To investigate how histone and RNA binding integrate, we utilized NMR spectroscopy. Specifically binding of ^15^N-labeled BD2 or BD4 to an H3K14ac peptide was assessed in the absence of, or when pre-bound to dsRNAI.

As expected, addition of H3K14ac peptide led to significant CSPs in BD2 and BD4 (Figure 4a,b (top panels), Supplementary Figure 9a, Supplementary Figure 10a). Consistent with previous studies, perturbations were most significant for residues in the ZA loop, BC loop, and residues just in the top of the αB and αC helices (Figure 4a,b – top panels). Comparison of CSPs induced by H3K14ac with those induced by dsRNAI (Figure 4a,b – top and middle panels) revealed that the histone and RNA binding pockets are partially overlapping for both BD2 and BD4. Residues in BD2 that are significantly perturbed by the addition of both dsRNAI and H3K14ac include E264, G266 and Q268 in the BC loop. For BD4 residues in the ZA loop are perturbed by both dsRNAI and H3K14ac (Supplementary Figure 11).

**Figure 4.**
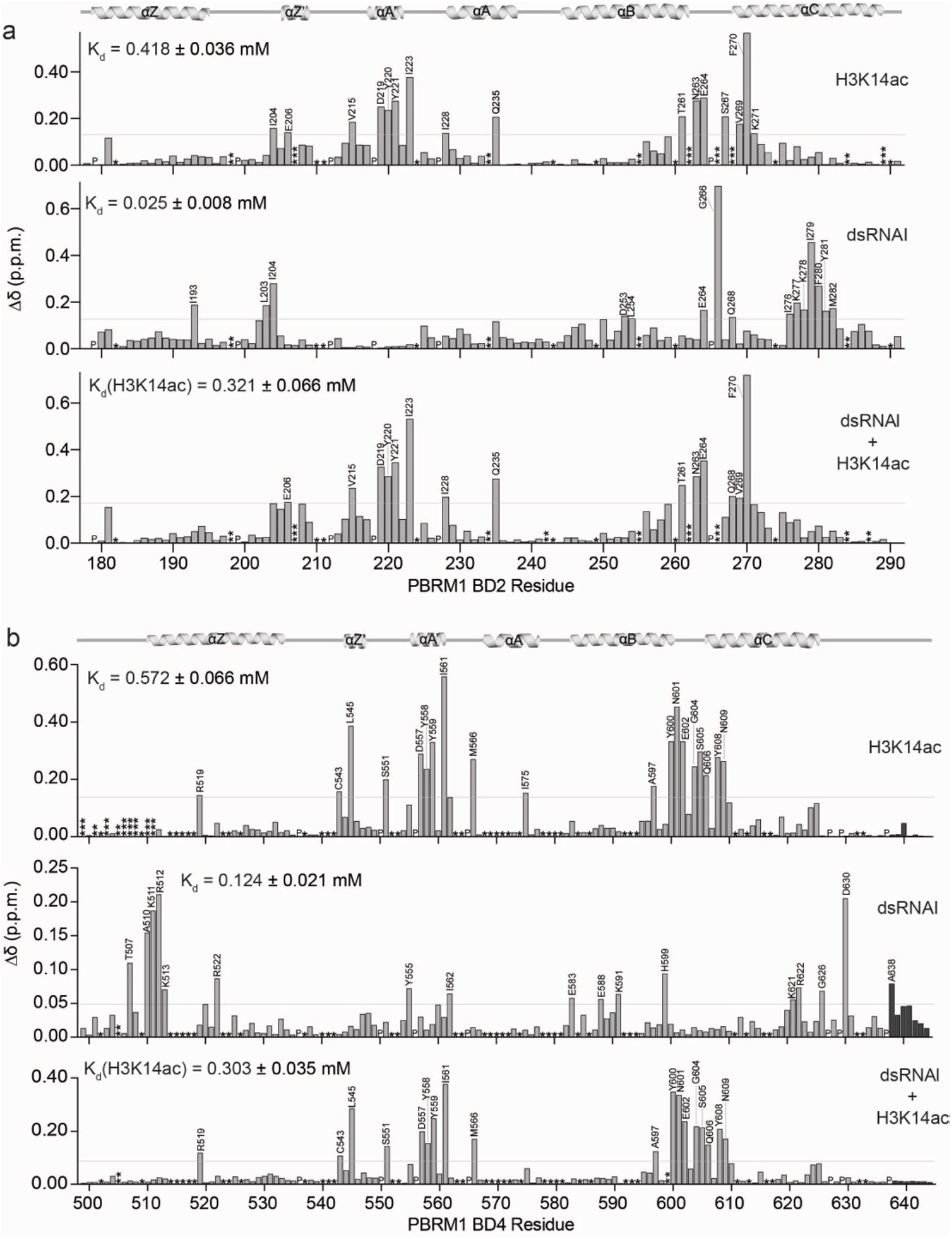
BD2 and BD4 have partially overlapping nucleic acid and histone binding pockets. **(a**,**b)** Normalized chemical shift changes (Δδ) between apo and H3K14ac-bound (top), dsRNAI-bound (middle), dsRNAI and H3K14ac-bound (bottom) for BD2 (a) and BD4 (b). Protein:dsRNAI, protein:H3K14ac, protein:dsRNAI:H3K14ac are in molar ratio of 1:6, 1:10 and 1:6:10 respectively for BD2. Protein:dsRNAI, protein:H3K14ac, protein:dsRNAI:H3K14ac are in molar ratio of 1:10, 1:40 and 1:10:15 respectively for BD4. Residues that are unassigned, merged with the addition of ligand, or broaden beyond detection upon addition of ligand, are marked as (*), (**) and (***) respectively. The secondary structure of the BD2 is denoted above the plots, and residues that were perturbed greater than the average plus two standard deviations (grey line) after trimming off the top 10% of CSPs are labelled.

To test if the BDs can bind to both histone tail and RNA simultaneously, H3K14ac peptide was titrated into dsRNAI-bound BD. For residues that are perturbed by peptide but not RNA, addition of peptide in the presence of RNA led to CSPs that are nearly identical to those seen without RNA present (Figure 4a,b – bottom panels, Supplementary Figure 9b, Supplementary Figure 10b). For residues that are perturbed by RNA but not peptide, addition of peptide in the presence of RNA did not lead to any further perturbations. Together this supports that BD2 and BD4 can bind to both RNA and peptide simultaneously. Notably, a subset of residues adopted a unique bound-state chemical shift in the presence of both RNA and peptide indicating a unique conformation of these residues in the ternary complex (see Supplementary Figure 12). For BD2, this includes; I204 in the ZA loop; I228, Q235 in the αA helix; Y261, N263, E264, S267, V269 in the BC loop; and K270, K271, I276, K277, K278, I279, F280, and M282 in the αC helix. For BD4 this includes; Y555, I562 in the ZA loop; E583 and Y599 in the αB helix; and K621 in the αC helix.

To assess the effect of RNA association on histone binding, affinities for the peptide were determined from the CSPs and compared in the absence and presence of RNA. For BD2 the K_d_ decreased from 0.41 mM ± 0.03 mM for H3K14ac alone to 0.32 mM ± 0.06 mM when pre-bound to dsRNAI (p-value <0.0001). Similarly, for BD4 the K_d_ for H3K14ac peptide decreased from 0.57 mM ± 0.06 mM to 0.30 mM ± 0.03 mM in the presence of RNA. Notably, this increase in affinity was not observed for BD4 when pre-bound to DNA (K_d_ = 0.63 mM ± 0.18 mM), revealing that this is unique to dsRNA.

### BD2 and BD4 nucleic acid binding contributes to PBRM1-mediated chromatin association and cellular function

To assess the functional importance of RNA binding, we first identified BD2 and BD4 mutants that inhibited RNA binding. From the NMR titrations with dsRNAI, residues with substantial CSPs upon titration of RNA (and not histone) were mutated to alanine: S275, K277 and Y281 in BD2, and K511, R512 and K513 in BD4. NMR ^1^H,^15^N-HSQC spectra confirmed that mutated BD2 and BD4 both retain their fold (Supplementary Figure 15). Titration of dsRNAI into BD2 S275A,K277A,Y281A revealed K_d_ = 0.20 mM ± 0.07 mM corresponding to a ∼8-fold reduction in affinity (Figure 5a, Supplementary Figure 15a). Similarly, titration dsRNAI into BD4 K511A,R512A,K513A revealed a K_d_ of 1.02 mM ± 0.04 mM corresponding to a ∼8-fold reduction in affinity (Figure 5a; Supplementary Figure 15b).

**Figure 5.**
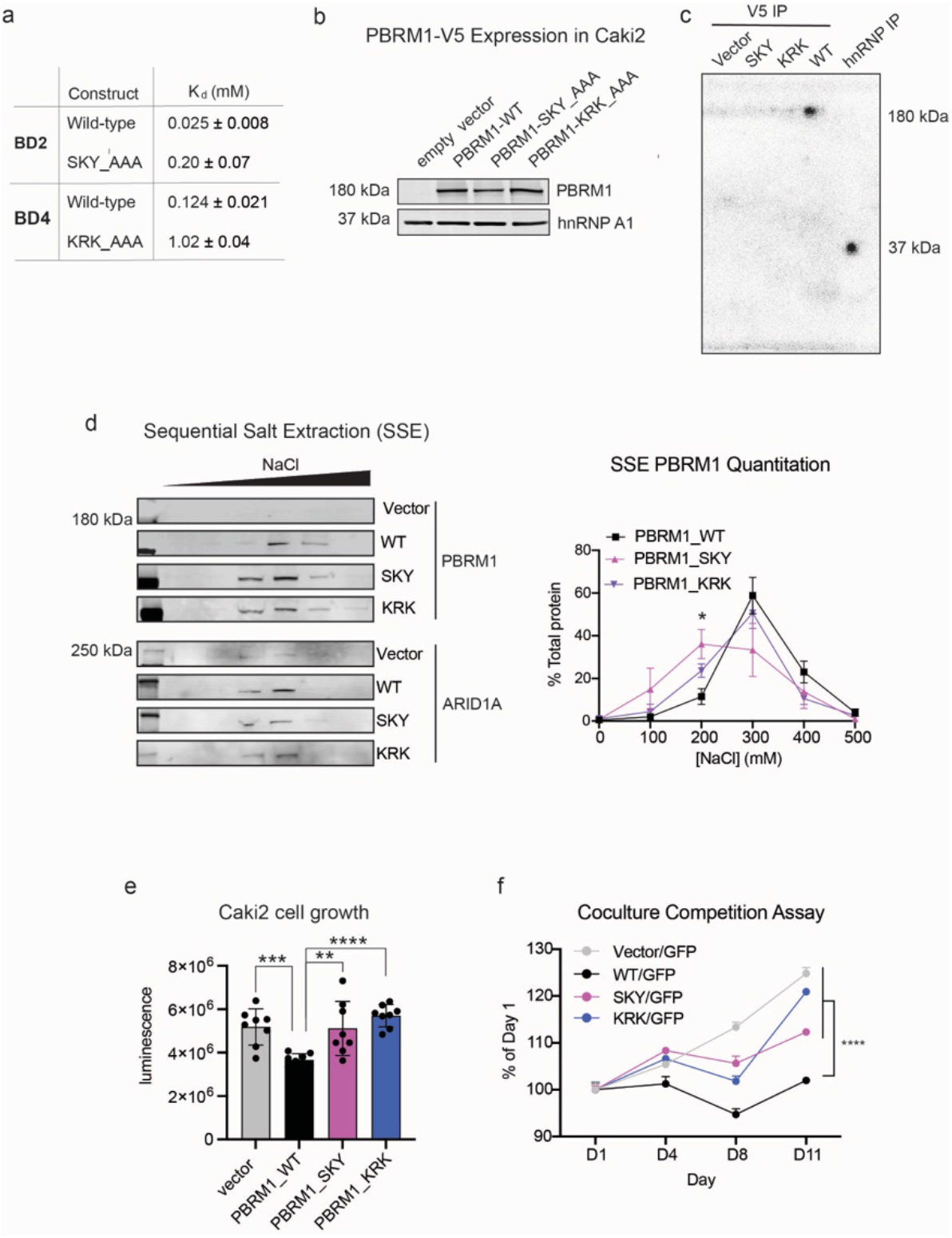
Mutation of RNA-binding residues disrupts PBRM1 function in cells. **(a)** Dissociation constants (K_d_) determined by NMR for BD2 and BD4 wild-type and mutant. **(b)** Immunoblot analysis of lysates from Caki2 renal clear cell carcinoma cells expressing full length WT PBRM1 or PBRM1 containing triple mutants in BD2 and BD4. hhnRNPA1 is included as a loading control. (**c)** Crosslinking immunoprecipitation (CLIP) performed in Caki2 cells. Phosphorimage of 32P-labeled immunoprecipitations of exogenous V5 tagged PBRM1, as well as positive control hnRNPA1 from UV crosslinked Caki2 cells. (**d)** (left) Representative immunoblot of PBRM1 (PBAF) and ARID1A (cBAF) elution by sequential salt extraction (SSE) in Caki2 cells with WT PBRM1 and PBRM1 mutants. (right) Analysis of binding affinity to chromatin by SSE of PBRM1 in the BD mutants indicated by the percentage of PBRM1 eluted at increasing NaCl concentrations. n = 4 independent biological replicates. A designation of * = p <0.05 (paired Student *t*-test). Error bars represent s.d. (**e)**. CellTiter-Glo® measurement of viable cells for seven days of culture of 2,000 cells plated in 96-well plates. A designation of * = p <0.05, ** = p <0.01, *** = p<0.001, **** = p<0.0001 (paired Student *t*-test). Error bars represent s.d. n = 8. (**f)** The change in the proportion of GFP negative cells compared to GFP positive cells as measured by flow cytometry. Equal numbers of GFP-labeled Caki2 cells and Caki2 cells expressing inducible PBRM1 were plated on day 0 and cells were harvested on designated time points for analysis. A designation of **** = p<0.0001 (paired Student *t*-test). Error bars represent s.d. n = 4.

The triple mutants in each BD (S275A,K277A,Y281A-SKY, and K511A,R512A,K513A-KRK) were incorporated into full length PBRM1 and re-expressed in the PBRM1-null renal cancer cell line Caki2 (Figure 5b) as previously described(42,61). To determine whether full-length PBRM1 directly associates with RNA in cells, we performed a crosslinking immunoprecipitation (CLIP) assay with ^32^P radiolabeled RNA. As a positive control, immunoprecipitation of hnRNP a 37 kDa RNA-binding protein, was included. Immunoprecipitation of the exogenously expressed PBRM1 under denaturing conditions with an antibody against V5 (Supplementary Figure 16a) reveals specific labelling of RNA at the same molecular weight only in cells expressing PBRM1 (Figure 5c, Supplementary Figure 16b). In comparison, PBRM1 with mutations in BD2 (SKY) or BD4 (KRK) have significantly reduced ^32^P labeling upon IP, indicating that the mutation decrease association with RNA in cells (Figure 5c, Supplementary Figure 16b).

Since BD2 and BD4 can associate with both DNA and RNA substrates, we next determined how mutations in BD2 and BD4 affect the overall affinity of PBRM1 for bulk chromatin containing RNA, DNA, histones, and associated proteins. For this, we employed a sequential salt extraction (SSE) assay, which measures the relative affinity of nuclear proteins to bulk chromatin(58). In Caki2 cells without PBRM1 expression, we previously found that PBAF subunits, such as ARID2, elute with a similar salt concentration as cBAF subunits such as ARID1A(42). Upon exogenous expression of PBRM1, all PBAF complex subunits elute with higher NaCl concentrations than cBAF subunit ARID1A, indicating that a subset of PBAF chromatin binding affinity is dependent on PBRM1. Here we observe a similar increase in the salt concentration required to elute WT PBRM1 compared to ARID1A (Figure 5d), but find that PBRM1 mutants SKY and KRK both elute with lower salt concentrations as compared to WT PBRM1. Notably, this loss in binding affinity is similar to what was previously observed with PBRM1 BD2 and BD4 bromodomain acetyl-lysine binding mutants(42). A decrease in binding affinity for the nucleic acid-binding mutants was consistent over multiple replicates, with a statistically significant difference in the percent of mutant PBRM1 eluted at 200 mM NaCl, compared to WT PBRM1 (Figure 5d).

Re-expression of wild type PBRM1 into Caki2 cells significantly slows growth compared to control(30), and this growth suppression is significantly reduced in the PBRM1 BD2 (SKY) and BD4 (KRK) mutant cells (Figure 5e, Supplementary Figure 16c) Due to the moderate effects of PBRM1 expression on Caki2 cell growth, as well as high variability in Caki2 growth rates, we developed a second FACS-based competition assay using GFP-labeled Caki2 cells. Using this assay, we found Caki2 GFP-labeled cells can outcompete Caki2 cells expressing WT PBRM1, but not Caki2 cells with vector alone (Figure 5f, Supplementary Figure 16d). Caki2 cells expressing PBRM1 with triple mutations in BD2 (SKY) and BD4 (KRK) display a greater ability to out compete Caki2-GFP labeled cells than Caki2 cells expressing WT PBRM1 (Figure 5f), which was consistent over multiple experiments (Supplementary Figure 16e). This indicates that nucleic acid binding through BD2 and BD4 contributes to the growth suppressive function of PBRM1 in renal cancer cells.

## Discussion

In this study, we show that PBRM1 BD2-BD5 bind to nucleic acids. We find that BD2 and BD4 preferentially bind to RNA over DNA *in vitro*, selectively associating with double-stranded RNA elements. The overall affinity and selectivity of BD2 is higher as compared to BD4. Notably, for both BDs the interaction with RNA leads to an enhanced affinity for acetylated histone tails.

In recent years, a number of known histone reader domains have been identified to also associate with DNA and/or RNA(62). This includes a subset of BDs, namely BRDT BD1 with DNA, BRG1/BRM BD with DNA, and BRD4 BD1 and BD2 with RNA(45–48). The molecular mechanism of association has been explored for the BRDT BD1(45) and the BRG1/BRM BD(46,48) while the mechanism of BRD4 BD association with RNA is not yet known. For the BRG1/BRM BD the binding pocket spans the αA helix, ZA loop and very N-terminal end of the αZ helix(46,48) While less completely defined, the BRDT BD1 DNA binding pocket includes the αZ helix(45). Here we found that RNA binds BD4 in a region N-terminal to αZ as well as in αZ, and C-terminal to αC and in αC. On the other hand, the BD2 RNA binding pocket is mainly located in the αC and αB helices, as well as the BC and ZA loops. While DNA binds more weakly to both BD2 and BD4, the binding pockets are nearly identical to that seen for RNA. Thus, while the acetyl lysine binding pocket is well conserved amongst BDs, the nucleic acid binding pockets characterized to date vary in their sequence composition and positioning on the BD. However, all align along a face of the alpha helical bundle and are comprised of a charged patch rich in arginine and lysine.

Our observation that the RNA binding enhances the histone tail binding activity of the BDs is in line with a previous study on BRD4(47), where interaction of the BDs with an enhancer RNA were found to augment the pull-down of acetylated histone peptides. It is interesting that we did not observe an increase in affinity of BD4 for histone tails when bound to DNA, as BRG1/hBRM association with DNA was also not seen to enhance the affinity for histone tails. Though this suggests this could be an effect unique to BD association with RNA, more work is needed to fully understand this.

Our data reveal that mutation of the RNA binding pocket decreases affinity for chromatin and inhibition of renal carcinoma cell proliferation. SWI/SNF subunits have previously been shown to interact with long non-coding RNAs (lncRNAs) (63–75). Interaction with lncRNAs can have a number of functional consequences including inhibition of activity by acting as a decoy, promoting activity through chromatin targeting or allosteric activation, or facilitating complex stability (76). While our results are consistent with a targeting or stability model, further studies are needed to determine the mechanism by which the RNA binding activity of the PBRM1 BDs is functioning *in vivo*. It should also be noted that while RNA is selected for *in vitro* under conditions of equal size and concentration, the relative importance of RNA versus DNA association in cells is not yet clear, and is quite likely context dependent. Future studies are needed to understand the interplay between RNA, DNA and histone binding, and the cooperative action of all six BDs in PBAF activity.

## Acknowledgements

We would like to thank Drs. David N.M. Jones and Chris Ptak for assistance with NMR, Drs. Brian Wimberly and Quentin Vicens for discussions regarding RNA construct design. Thanks to the members of the Musselman laboratory as well as Drs. Nicholas Schnicker and Zhen Xu for valuable discussions regarding this project. Thanks to Dr. Aktan Alpsoy for assistance with cloning.

