## Supplementary Figures for "PBRM1 BD2 and BD4 associate with RNA to facilitate chromatin association"

a

```
BD1  ----------SMTV---------DPI-AVCHELYNTIRDYK-DEQGRLLCELFIRAPKRRNQPDYYEVVSQPI
BD2  ----------SMSP--------AY-LKEILEQLLEAIVVAT-NPSGRLISELFQKLPSKVQYPDYYAIIKEPI
BD3  ----------SM-------------------QLYDTVRSCR-NNQGQLIAEPFYHLPSKKKYPDYYQQIKMPI
BD4  MGDSMISSATSDTGSAKRKSK-KNIRKQRMKILFNVVLEAREPGSGRRLCDLFMVKPSKKDYPDYYKIILEPM
BD5  ----------SMSGISPKKSKYMTPMQQKLNEVYEAVKNYT-DKRGRRLSAIFLRLPSRSELPDYYLTIKKPM
BD6  ----------SMNV--------TLLIQELIHNLFVSVMSHQ-DDEGRCYSDSLAEIPAVD--PNF--PNKPPL
               *                     :   :        *:  .  :   *     *::      *:

BD1  DLMKIQQKLKMEEYDDVNLLTADFQLLFNNAKSYYKPDSPEYKAACKLWDLYLRTRNEFVQKG----------
BD2  DLKTIAQRIQNGSYKSIHAMAKDIDLLAKNAKTYNEPGSQVFKDANSIKKIFYMKKAEIEHH-----------
BD3  SLQQIRTKLKNQEYETLDHLECDLNLMFENAKRYNVPNSAIYKRVLKLQQVMQAKKKELARRDD---I-----
BD4  DLKIIEHNIRNDKYAGEEGMIEDMKLMFRNARHYNEEGSQVYNDAHILEKLLKEKRKELGPLPDDDDMASPA-
BD5  DMEKIRSHMMANKYQDIDSMVEDFVMMFNNACTYNEPESLIYKDALVLHKVLLETRRDLEG------------
BD6  TFDIIRKNVENNRYRRLDLFQEHMFEVLERARRMNRTDSEIYEDAVELQQFFIKIRDELCKNGEI--LLSPAL
      :  *  .:    *   . :  .:  : ..*       *  :: .  : ..    : ::

BD1  ----E---------------------
BD2  -------------------------E
BD3  ----E---------------------
BD4  --------------------------
BD5  ----D---------------------
BD6  SYTTKHLHNDVEKERKEKLPKEIEED
```

b

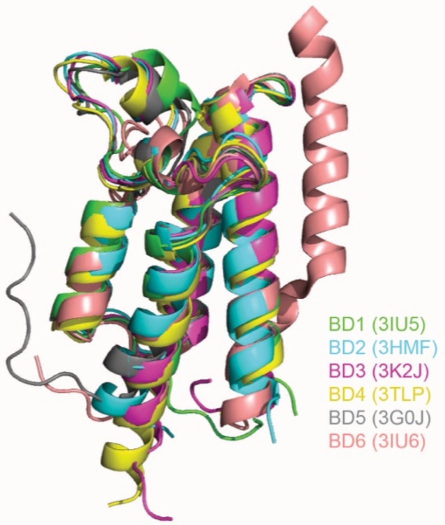


**SI Figure 1.** Sequence alignment (performed in T-Coffee) of the PBRM1 bromodomains (BD1-BD6). Included are the amino acids from the constructs used in this study. Residues are denoted as identical (* and shown in red), strongly similar (:), or weakly similar (.). Overlay of the previously solved crystal structures of BD1-BD6.

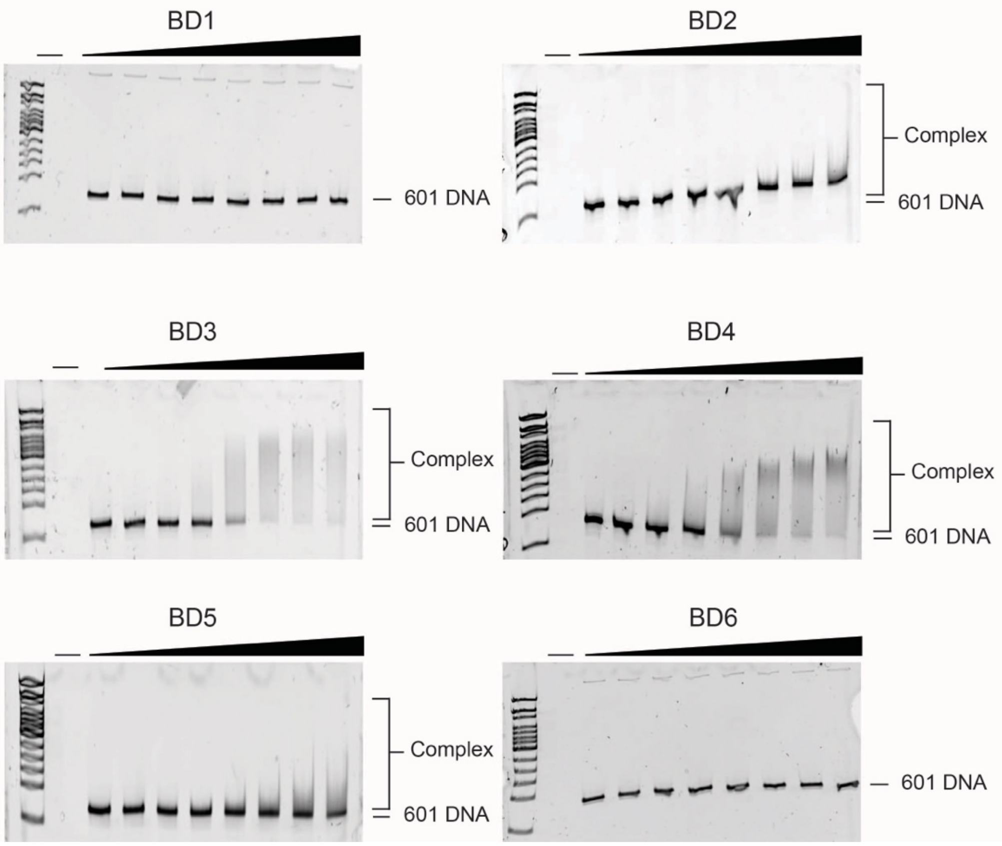


**SI Figure 2.** Electromobility shift assays (EMSAs) carried out with the Widom 601 DNA with individually purified BD1-BD6. Gels were run with a 100 bp DNA ladder on the left and stained with ethidium bromide for visualization.

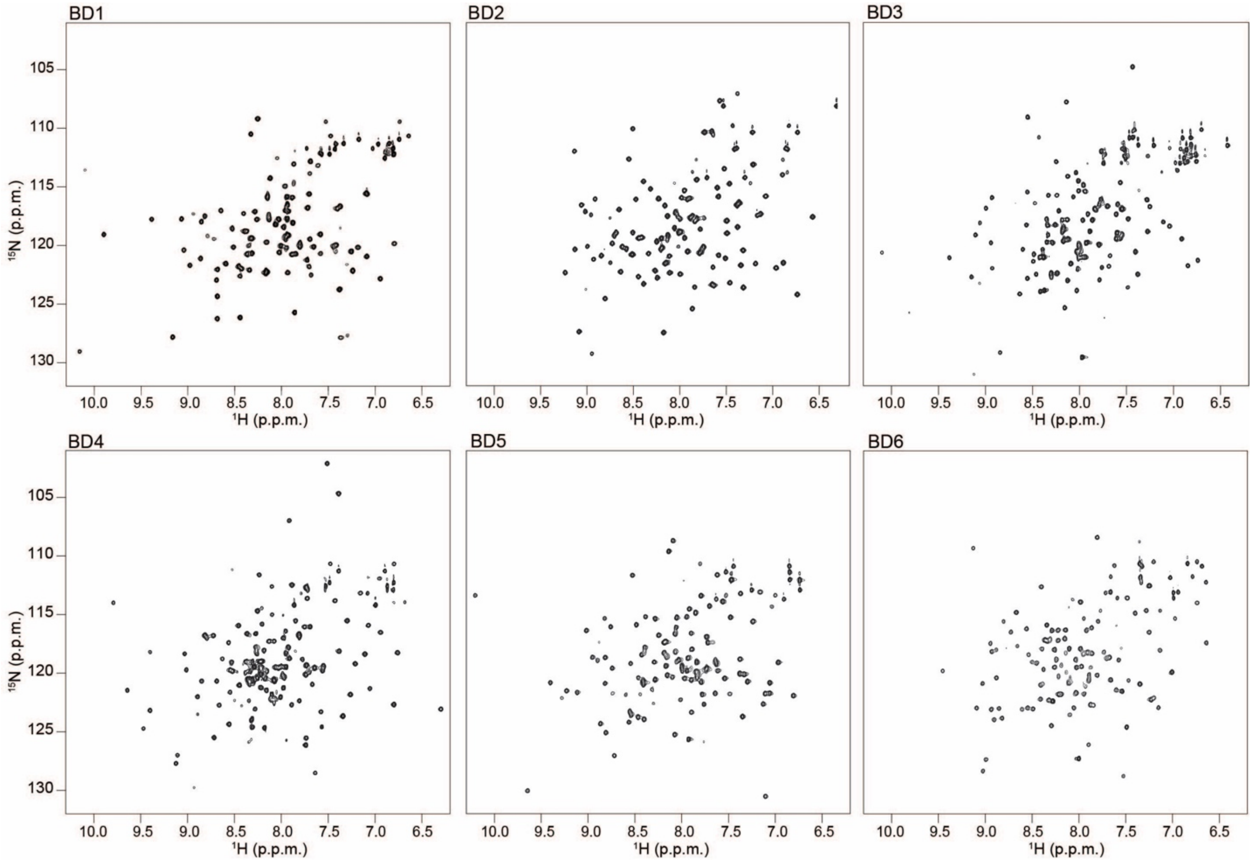


**SI Figure 3.** Individual $^{1}$H-$^{15}$N HSQC spectra of BD1-BD6. Spectra were collected on 0.1 mM in 93% $H_2O$/7% $D_2O$ samples in 800 MHz Bruker Avance II at 25 °C.

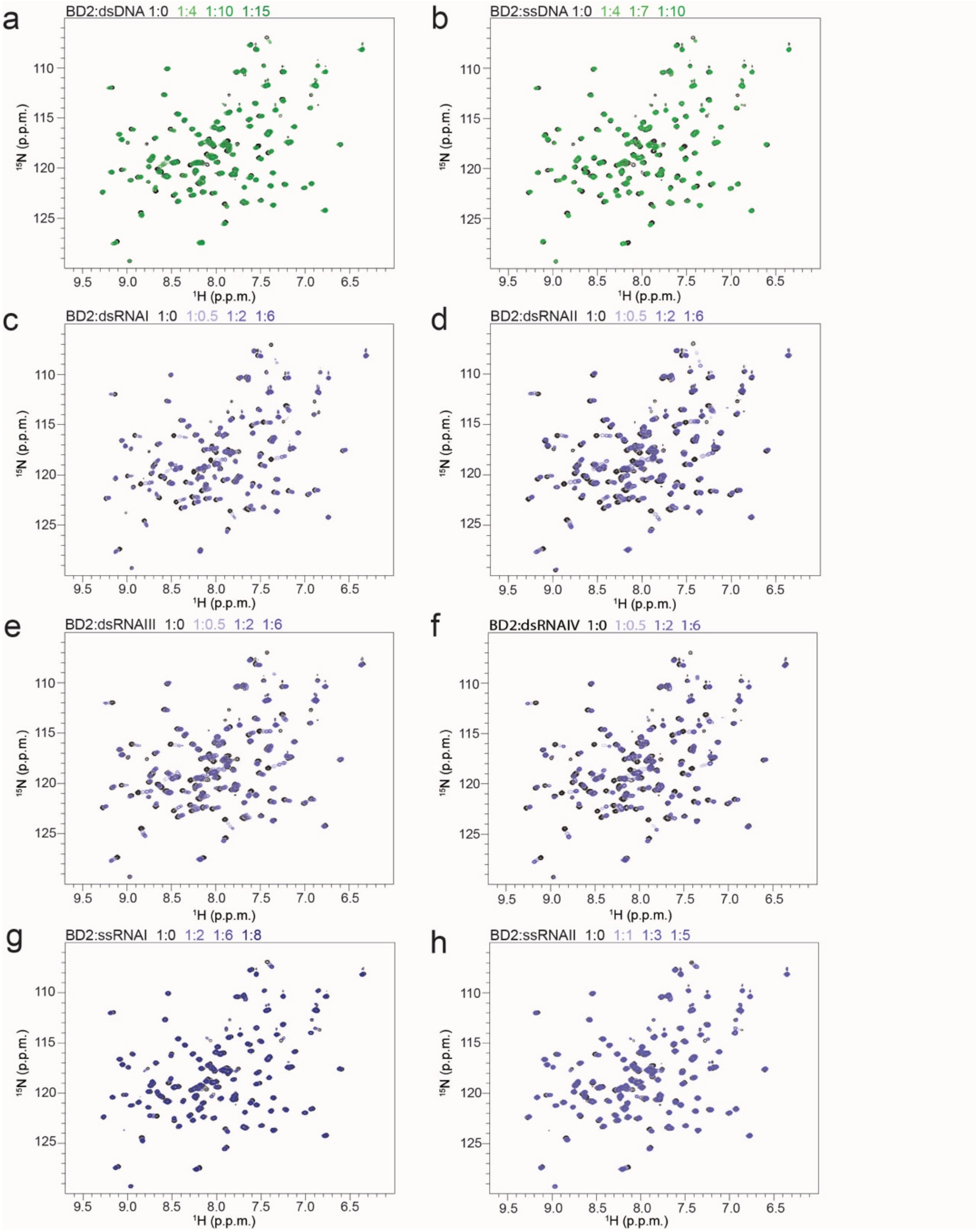


**SI Figure 4.** Overlay of $^{1}$H-$^{15}$N HSQC spectra of $^{15}$N-BD2 upon titration of (a) dsDNAI (b) ssDNA (c) dsRNAI (d) dsRNAII (e) dsRNAIII (f) dsRNAIV (g) ssRNAI (h) ssRNAII. Spectra are color coded according to protein:nucleic acid molar ratio as shown in the legend. dsDNA titration was collected at 1:0, 1:1, 1:2, 1:4, 1:6, 1:10, 1:12, 1:15, ssDNA was collected at 1:0, 1:0.5, 1:1, 1:2, 1:4, 1:7, 1:10, dsRNAI - dsRNAIV were collected at 1:0, 1:0.1, 1:0.25, 1:0.5, 1:1, 1:2, 1:4, 1:6, ssRNAI was collected at 1:0, 1:1, 1:2, 1:4, 1:6, 1:8 and ssRNAII was collected at 1:0, 1:0.5, 1:1, 1:2, 1:3, 1:4, 1:5. Protein concentration was 100 μM. For clarity, only 4 points are displayed.

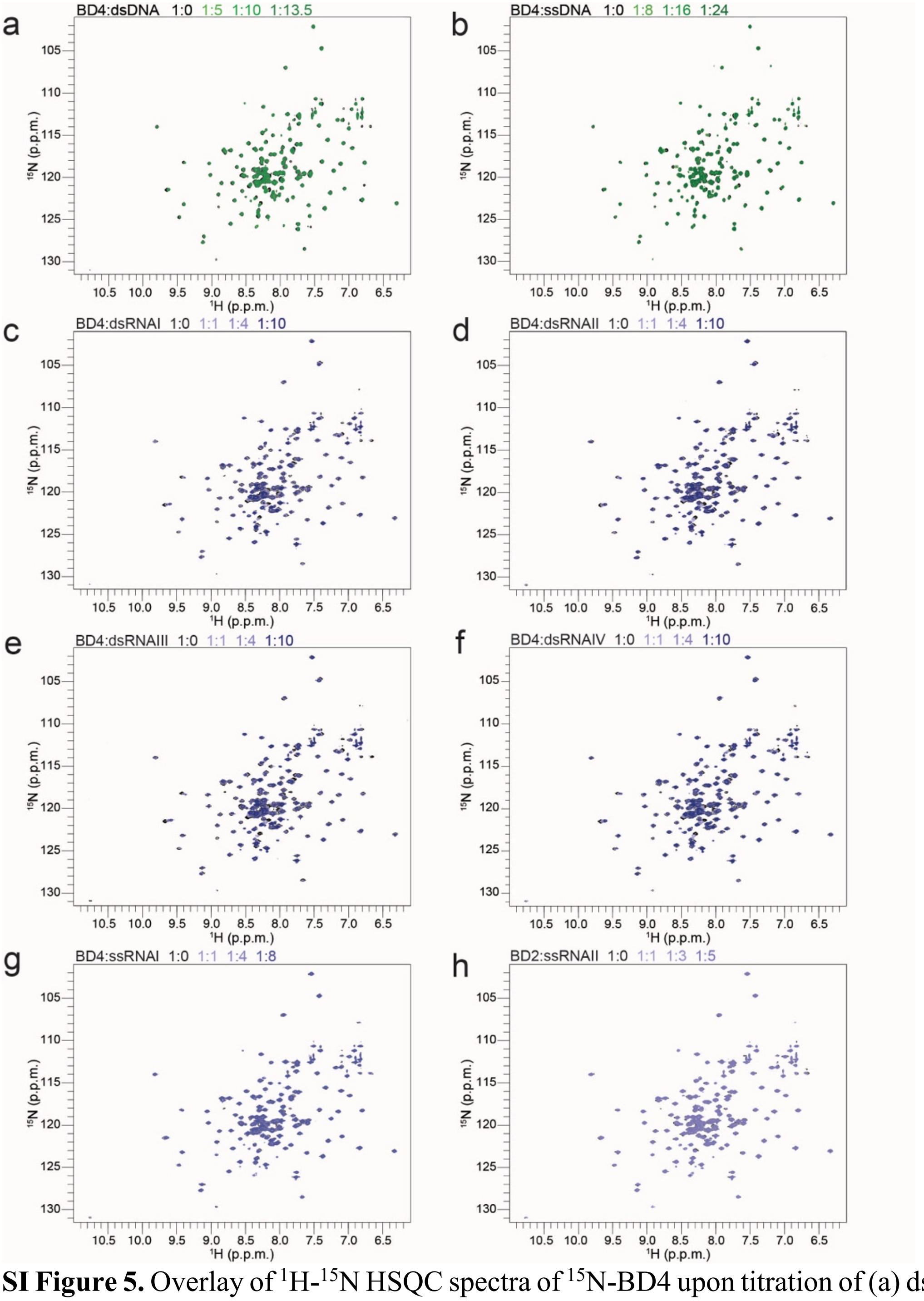


**SI Figure 5.** Overlay of $^{1}$H-$^{15}$N HSQC spectra of $^{15}$N-BD4 upon titration of (a) dsDNAI (b) ssDNA (c) dsRNAI (d) dsRNAII (e) dsRNAIII (f) dsRNAIV (g) ssRNAI (h) ssRNAII. Spectra are color coded according to protein:nucleic acid molar ratio as shown in the legend. dsDNA titration was collected at 1:0, 1:0.5, 1:1, 1:2, 1:2.5, 1:5, 1:7.5, 1:10, 1:13.5, ssDNA was collected at 1:0, 1:0.5, 1:1, 1:4, 1:8, 1:12, 1:16, 1:24, dsRNAI - dsRNAIV were collected at 1:0, 1:0.5, 1:1, 1:2, 1:3, 1:4, 1:6, 1:10, ssRNAI was collected at 1:0, 1:1, 1:2, 1:4, 1:6, 1:8 and ssRNAII was collected at 1:0, 1:0.5, 1:1, 1:2, 1:3, 1:4, 1:5. Protein concentration was 0.1 mM. For clarity, only 4 points are displayed.

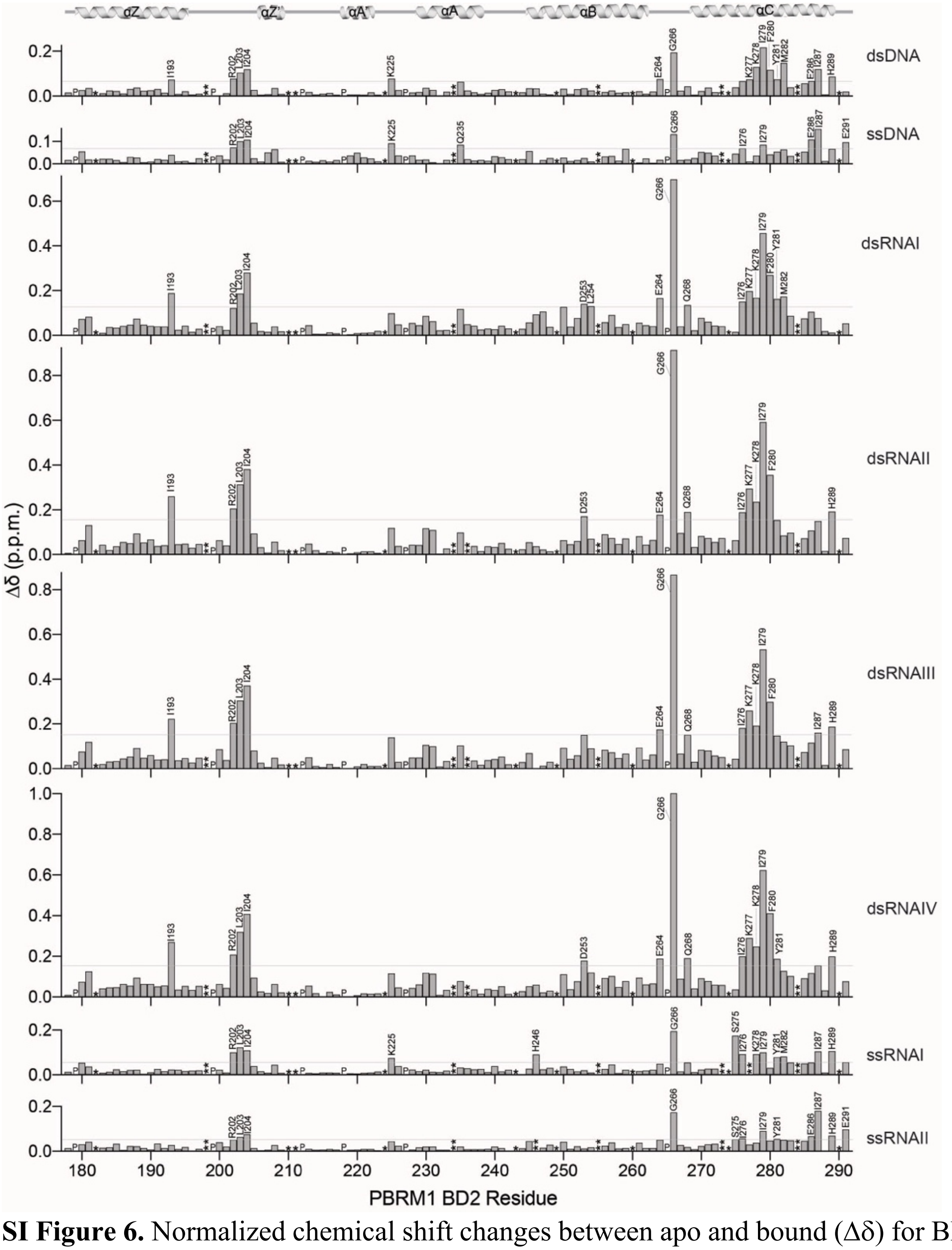


**SI Figure 6.** Normalized chemical shift changes between apo and bound (Δδ) for BD2 are plotted as a function of residue in the presence of dsDNAI, ssDNA, dsRNAI, dsRNAII, dsRNAIII, dsRNAIV, ssRNAI and ssRNAII (from top to bottom). Final molar ratios are 1:15, 1:10, 1:6, 1:6, 1:6, 1:6, 1:8 and 1:5, respectively. Residues that are unassigned or untrackable due to overlap are marked as (*) and (**) respectively. The secondary structure of the PBRM1 BD2 is denoted above the plots, and residues that were perturbed greater than the average plus two standard deviations after trimming off the top 10% are labelled. A grey line marks this level of significance for each titration.

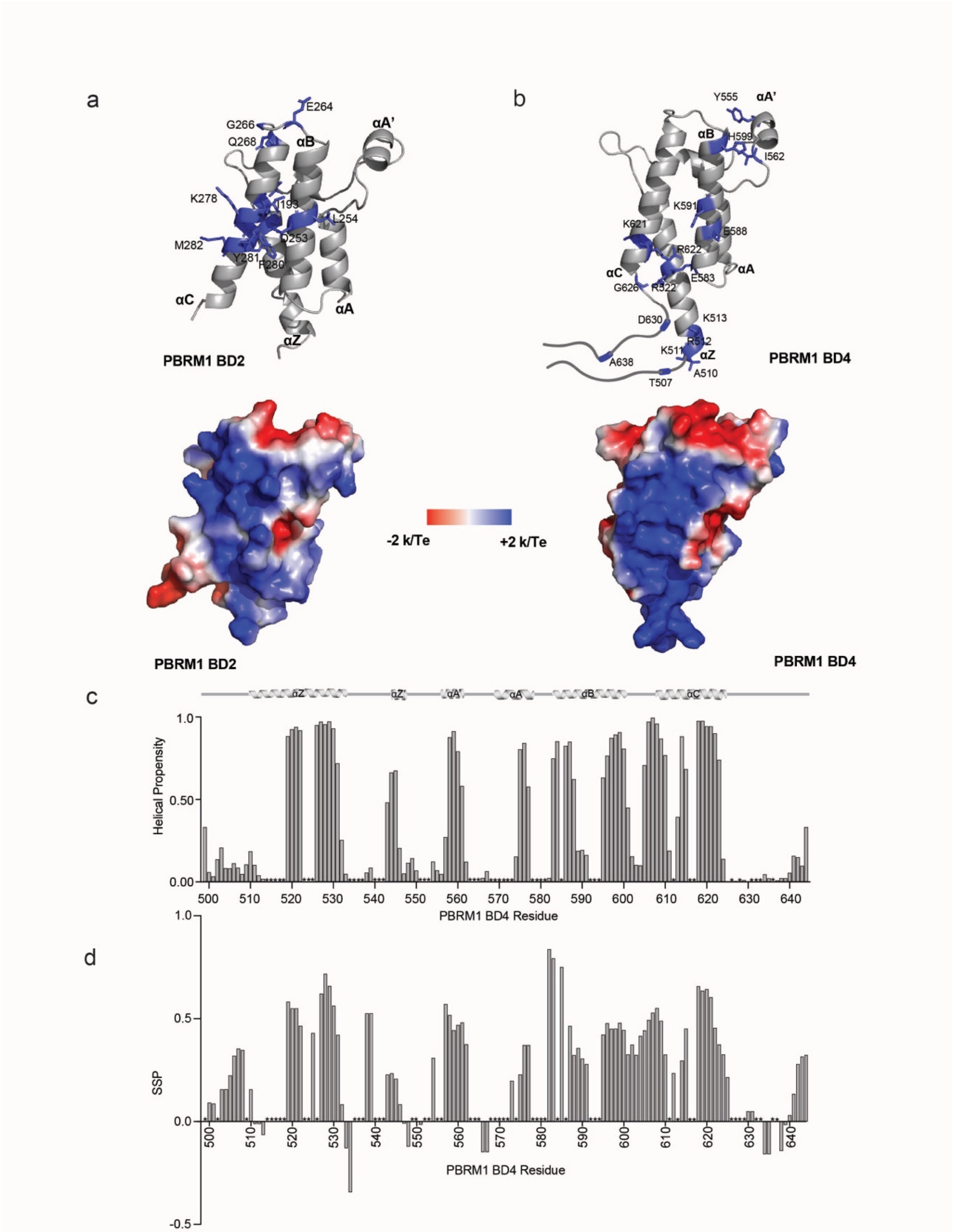


**SI figure 7.** Residues perturbed by addition of dsRNAI for (a) BD2 and (b) BD4 are highlighted as sticks and colored blue on the previously solved structures of each BD (PDB ID 3HMF and 3TLP). Below are the corresponding surface electrostatics calculated in pymol using the APBS plugin. (c) TALOS+ plot and (d) Secondary Structure Prediction (SSP) plot for BD4.

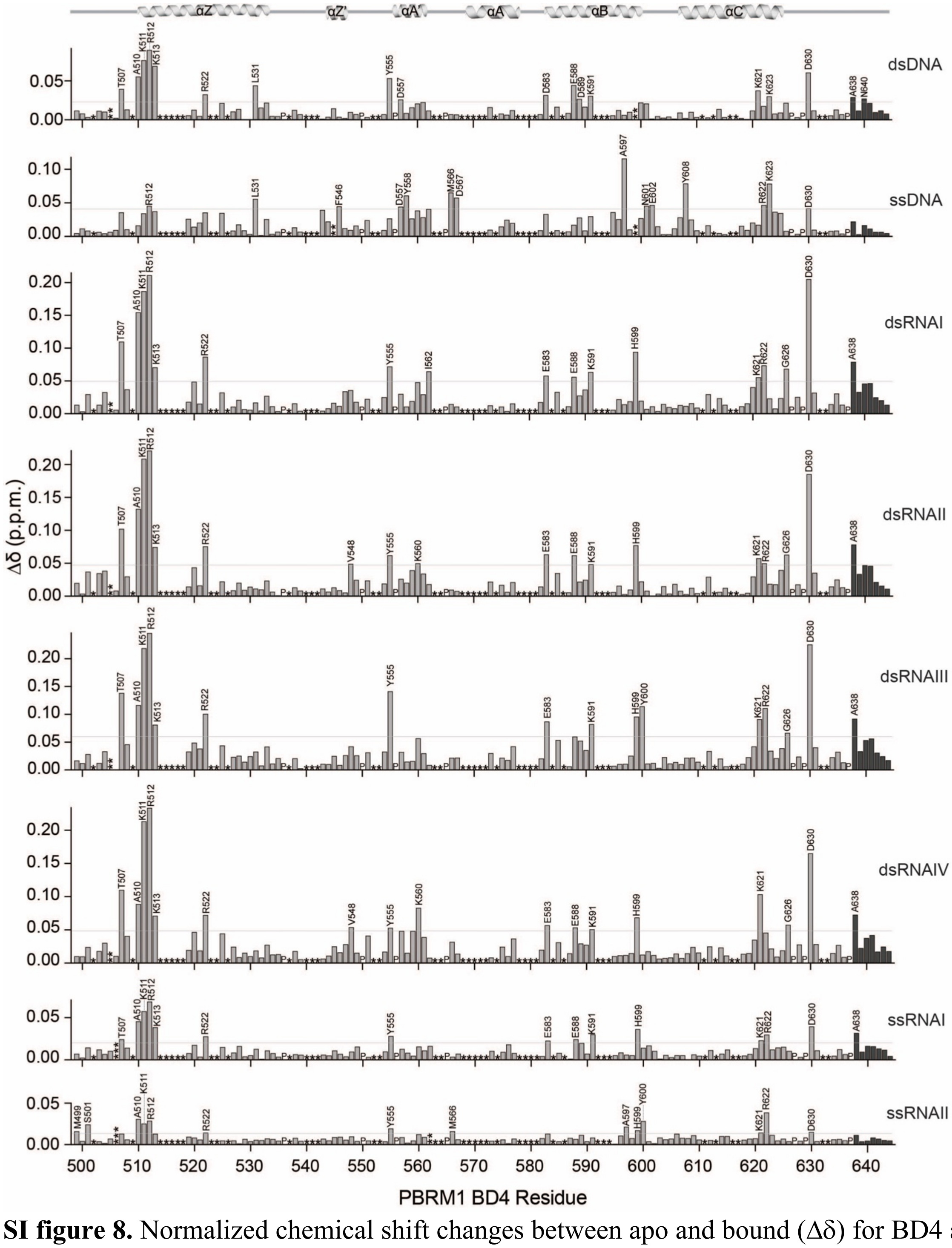


**SI figure 8.** Normalized chemical shift changes between apo and bound (Δδ) for BD4 are plotted as a function of residue upon titration of dsDNAI, ssDNA, dsRNAI, dsRNAII, dsRNAIII, dsRNAIV, ssRNAI and ssRNAII (from top to bottom). Final molar ratios are 1:13.5, 1:24, 1:10, 1:10, 1:10, 1:10, 1:8 and 1:5, respectively. Residues that are unassigned, untrackable due to overlap, or broaden beyond detection upon addition of ligand are marked as (*), (**) and (***) respectively. The secondary structure of the PBRM1 BD4 is denoted above the plots, and residues that were perturbed greater than the average plus two standard deviations after trimming off the top 10% of CSPs are labelled. A grey line marks this level of significance for each titration.

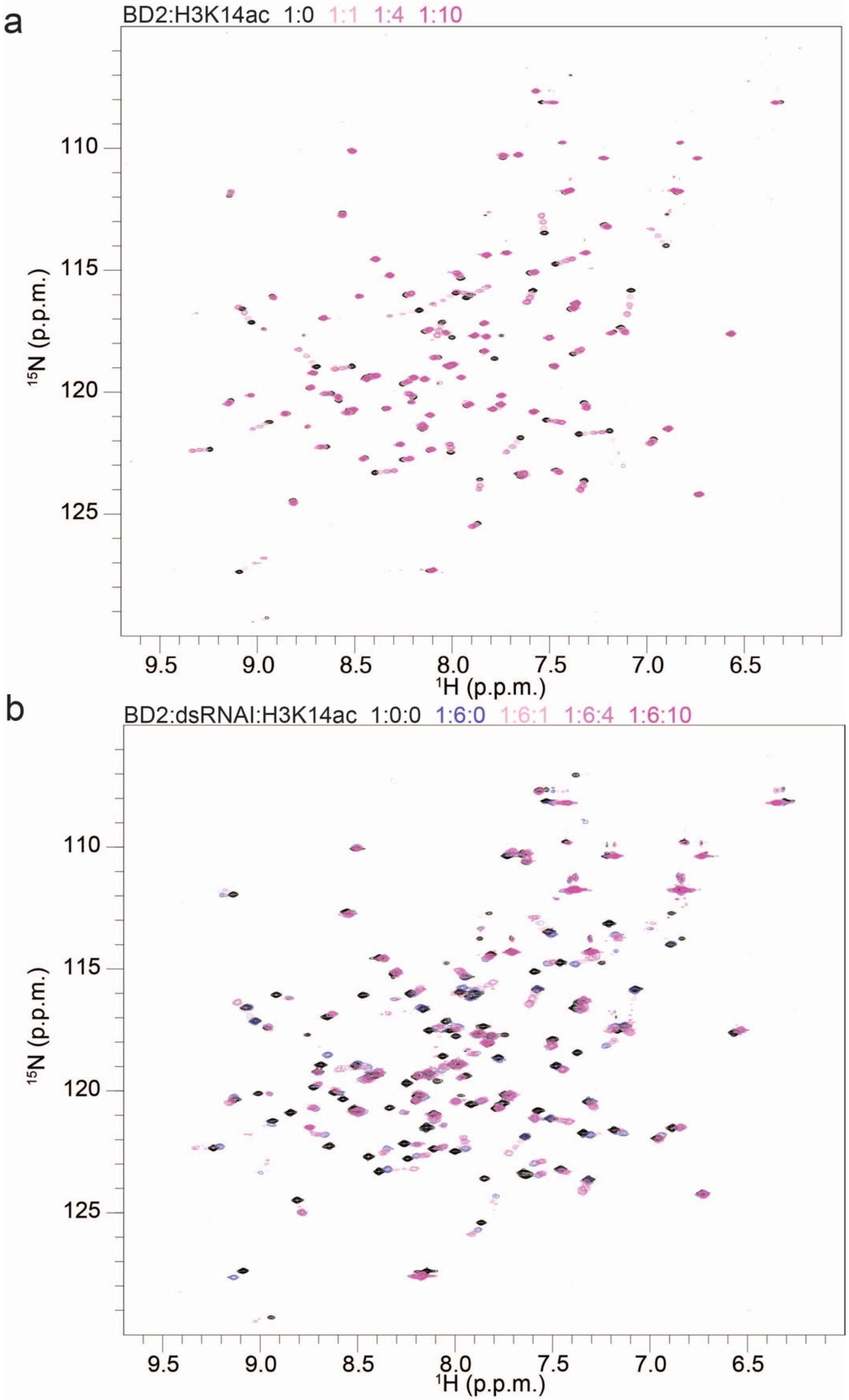


**SI figure 9.** Overlay of $^{1}H$-$^{15}N$ HSQC spectra of $^{15}N$-BD2 upon titration of (a) H3K14ac and (b) both dsRNAI and H3K14ac. Spectra are color coded according to legends.

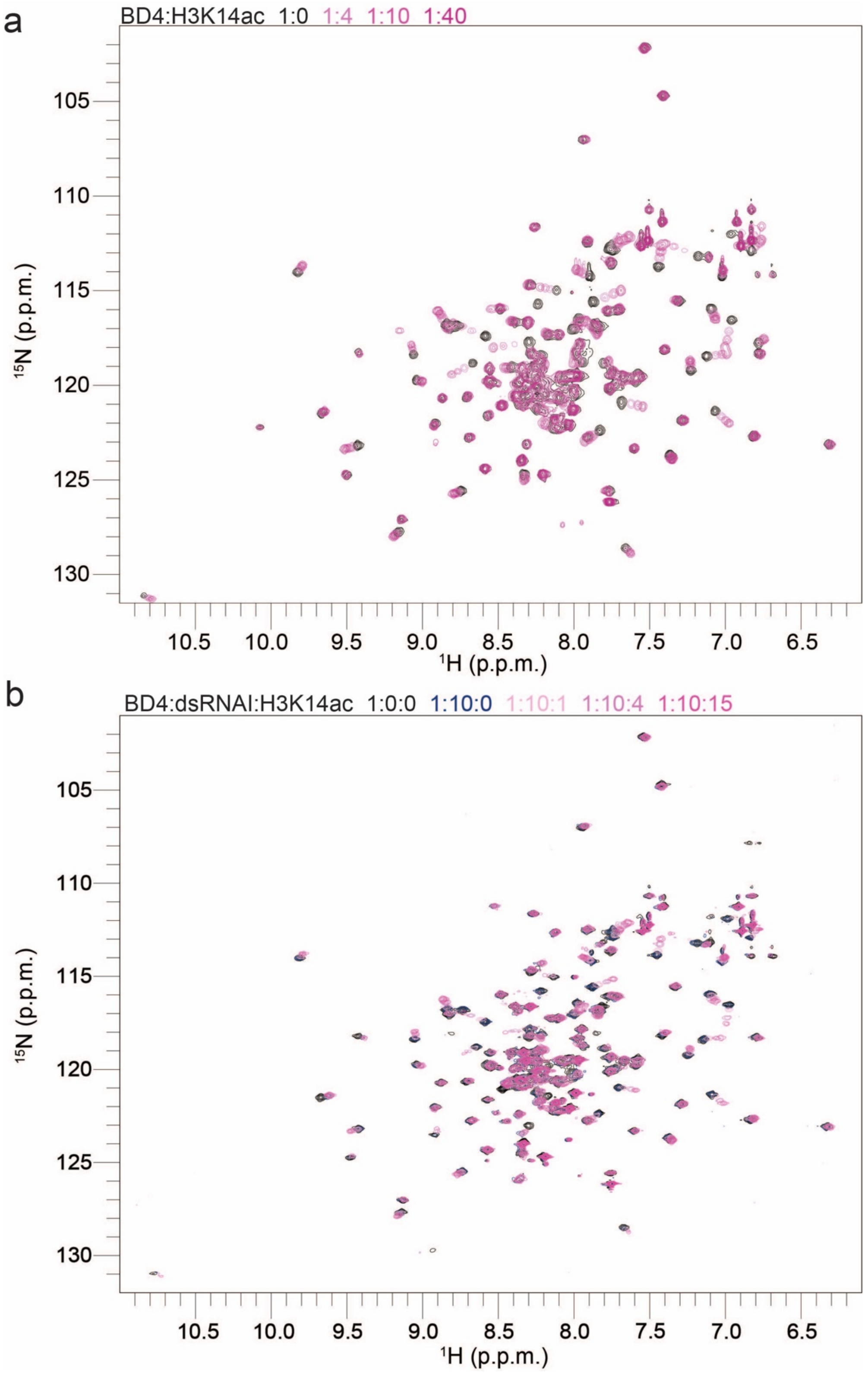


**SI figure 10.** Overlay of $^{1}H$-$^{15}N$ HSQC spectra of $^{15}N$-BD4 upon titration of (a) H3K14ac and (b) both dsRNAI and H3K14ac. Spectra are color coded according to legends.

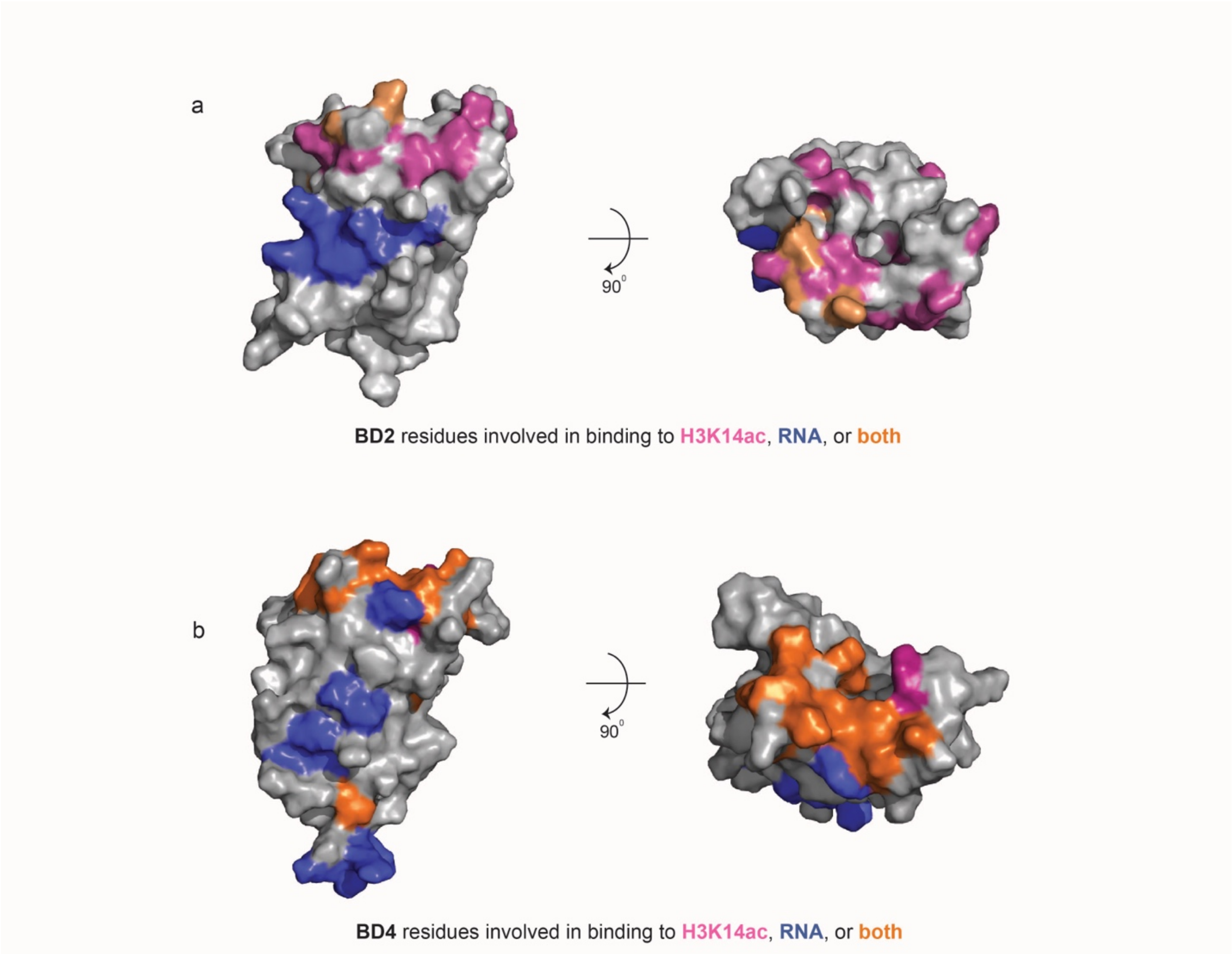


**SI figure 11.** Overlap in the residues that are perturbed upon binding H3K14ac and dsRNAI for BD2 (a) or BD4 (b). A surface representation of BD2 (PDB ID 3HMF) or BD4 (PDB ID 3TLF) with residues colored according to their involvement in binding to H3K14ac peptide (pink), dsRNAI (purple) or both (orange).

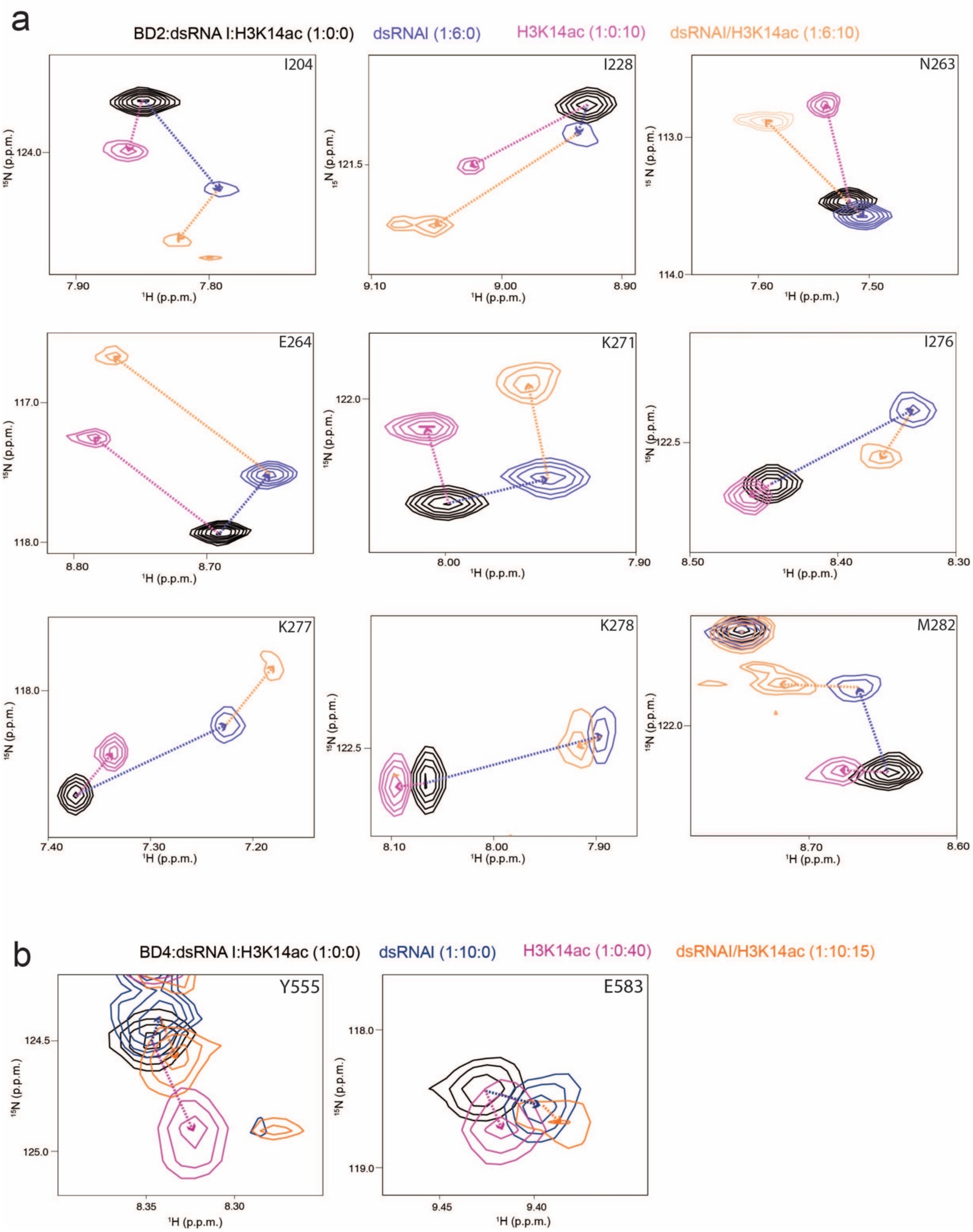


**SI figure 12**. Chemical shift trajectories for selected residues from $^{1}$H-$^{15}$N HSQC spectral overlays for $^{15}$N-BD2 (a) or $^{15}$N-BD4 (b). Shown are apo (black) in the presence of H3K14ac (pink), dsRNAI (purple), or both dsRNAI and H3K14ac (orange).

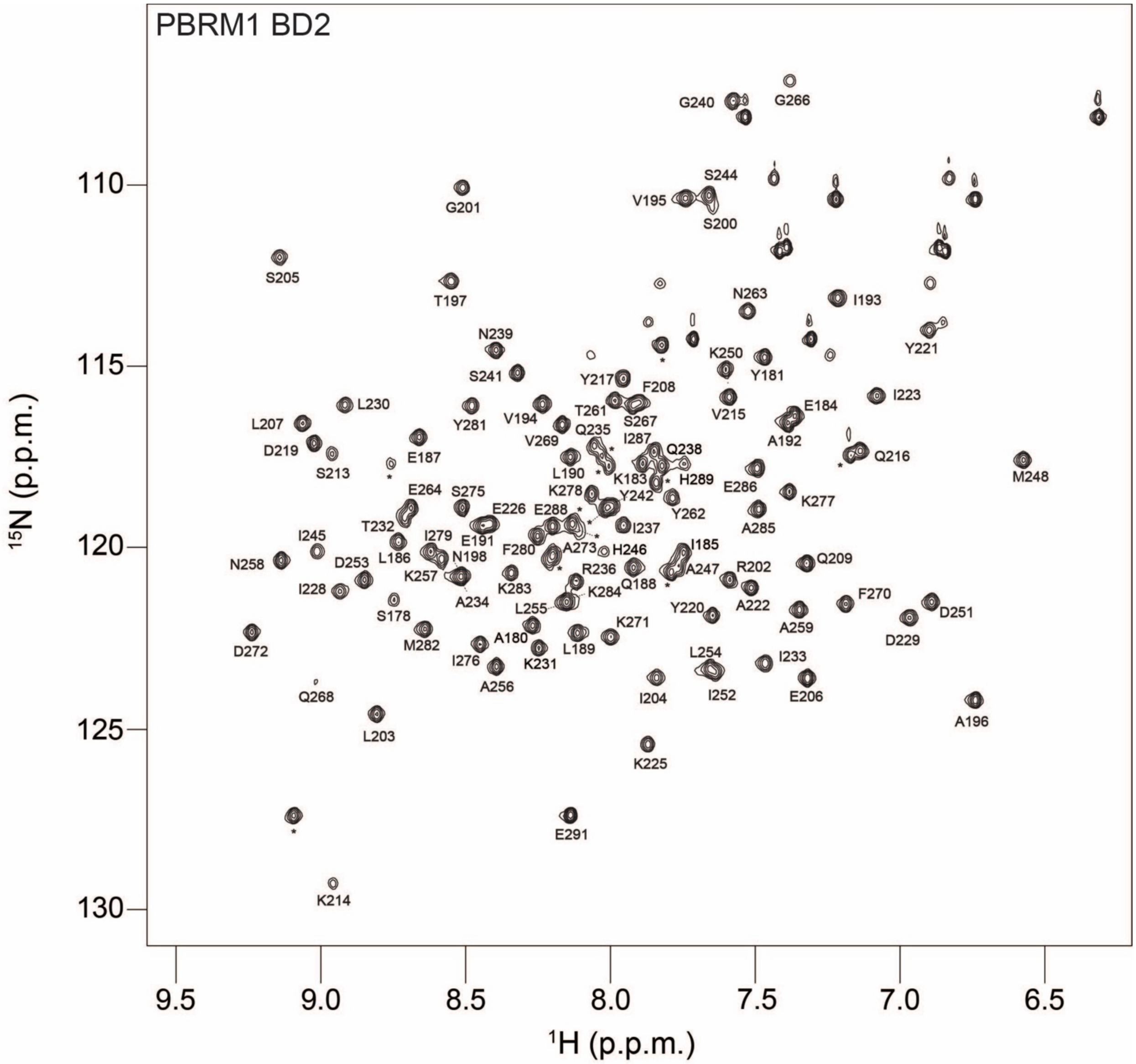


**SI figure 13.** Backbone assignments for BD2. The construct contains 116 residues, 6 of which are prolines. Thus, assuming fast exchange, 110 resonances would be expected. Of the 104 peaks observed 94 were successfully assigned.

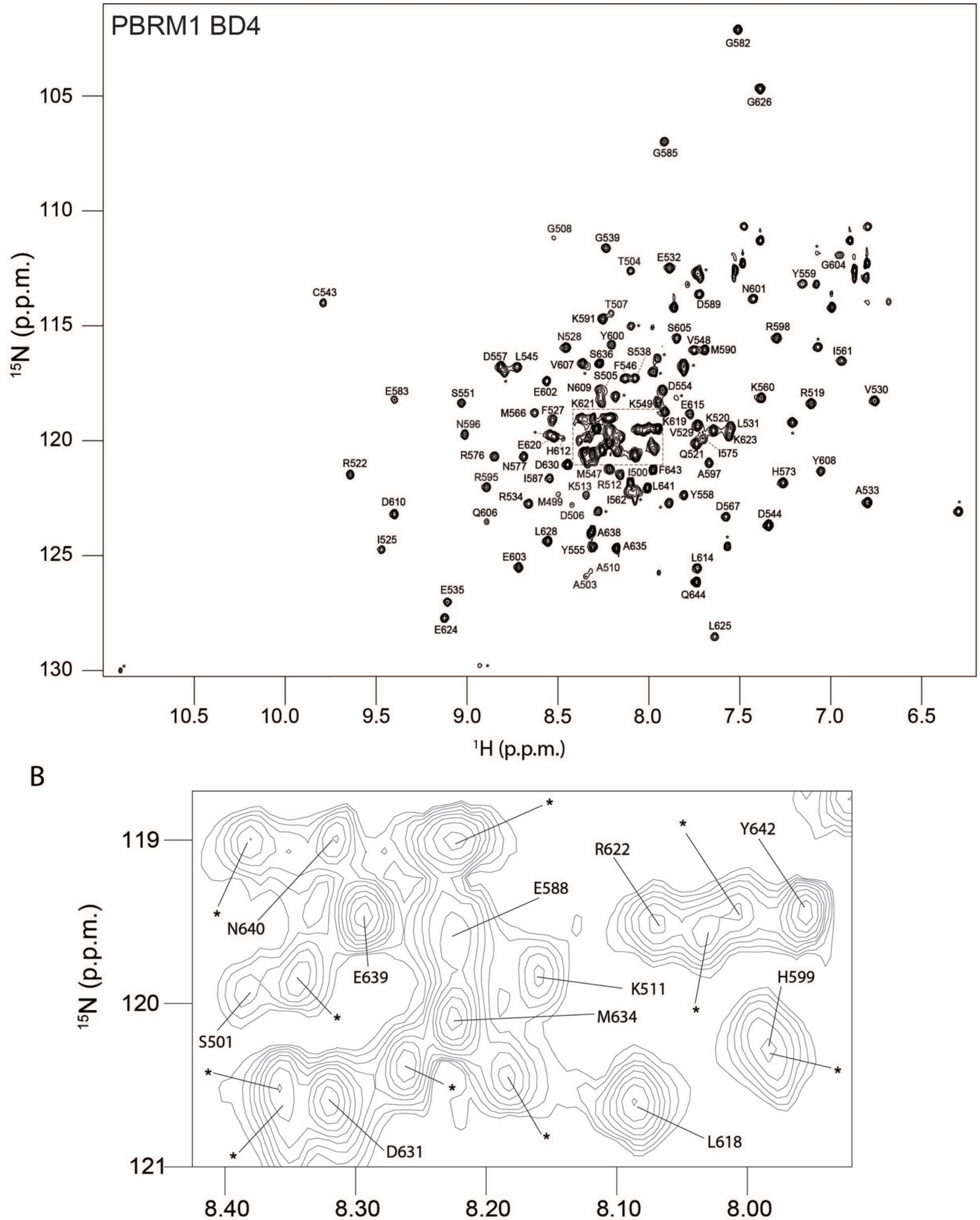


**SI figure 14.** Backbone assignments for BD4. The construct contains 150 residues, 7 of which are prolines. Thus, assuming fast exchange, 143 resonances would be expected. Of the 119 peaks observed 96 were successfully assigned.

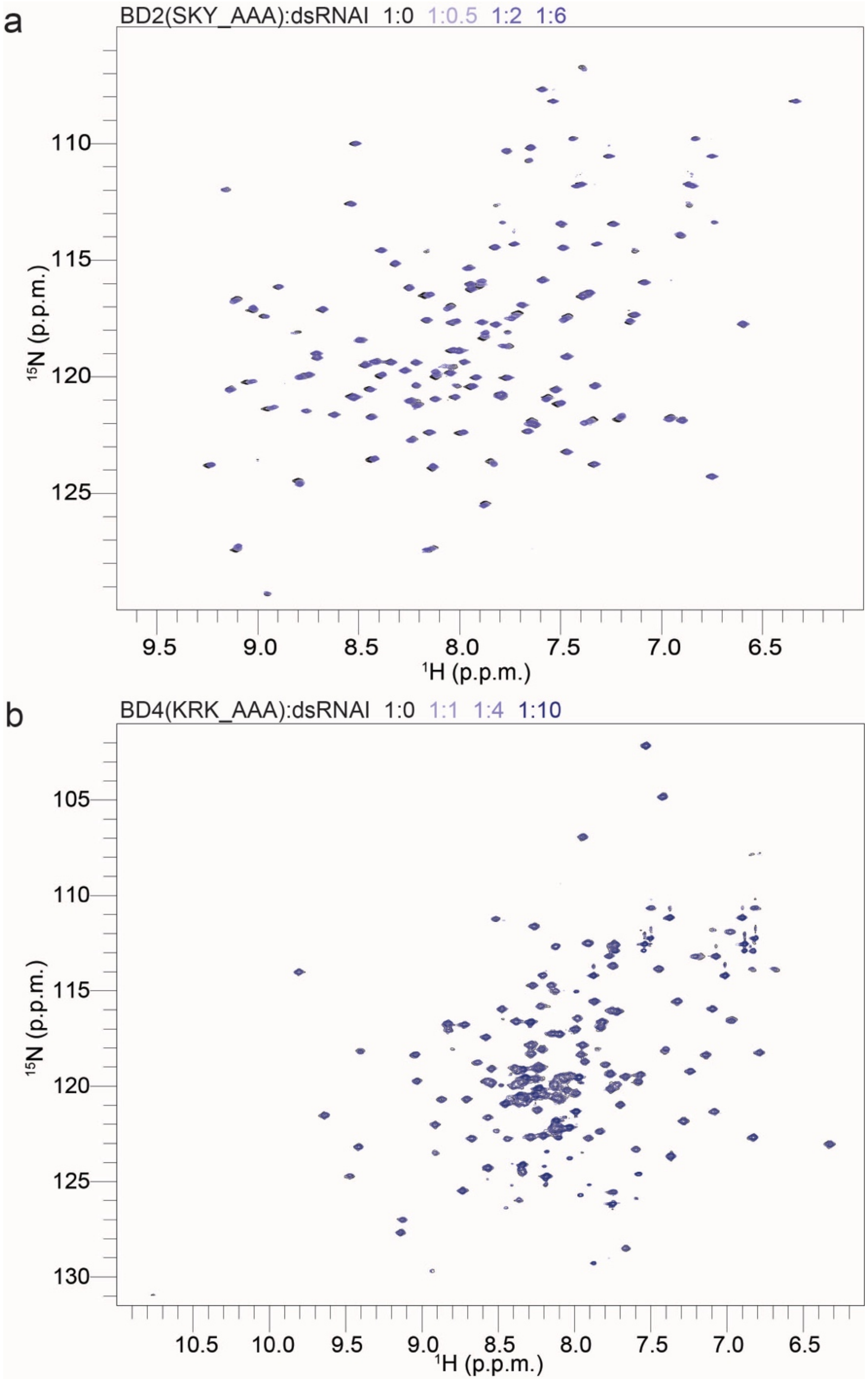


**SI figure 15.** Overlay of $^{1}$H-$^{15}$N HSQC spectra of (a) $^{15}$N-BD2 SKY_AAA or (b) $^{15}$N-BD4 KRK_AAA upon titration of dsRNAI. Spectra are color coded according to protein:dsRNAI molar ratio as shown in the legend. For BD2 spectra were collected at 1:0, 1:0.1, 1:0.25, 1:0.5, 1:1, 1:2, 1:4, 1:6. Protein concentration was 0.1 mM. For BD2 spectra were collected at 1:0, 1:0.5, 1:1, 1:2, 1:3, 1:4, 1:7, 1:10. Protein concentration was 0.1 mM. For clarity, only 4 points are displayed.

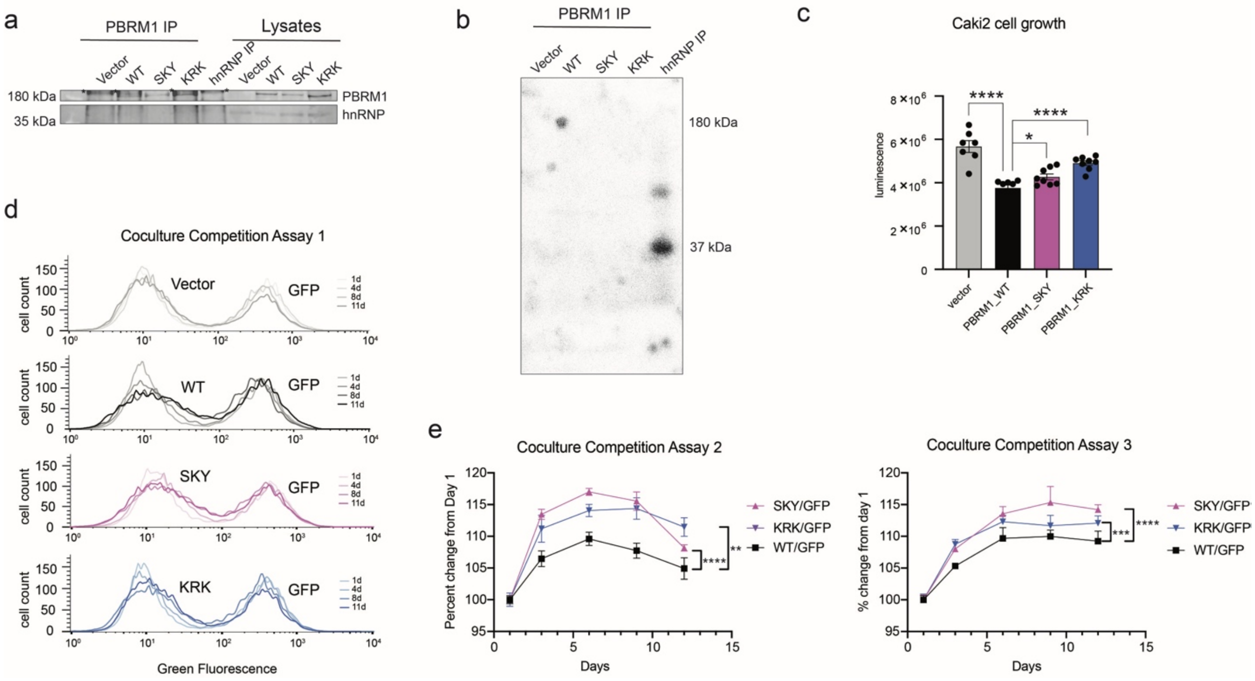


**SI Figure 16. (**a) Immunoblot of immunoprecipitation from UV crosslinked Caki2 cells (CLIP). * indicates a non-specific band. (b) Phosphorimage of 32P-labeled immunoprecipitations of exogenous V5 tagged PBRM1, as well as positive control hnRNPA1 from UV crosslinked Caki2 cells. (c) CellTiter-Glo® measurement of viable cells for seven days of culture of 2,000 cells plated in 96-well plates. A designation of * = p <0.05, ** = p <0.01, *** = p<0.001, **** = p<0.0001 (paired Student *t*-test). Error bars represent s.d. n = 8. (d) Representative histograms of GFP+Caki2 cells and GFP-Caki2 cells with the designated PBRM1 re-expression status. The same initial cell populations were assayed over 11d of culture. (e) The change in the proportion of GFP negative cells compared to GFP positive cells as measured by flow cytometry. Equal numbers of GFP-labeled Caki2 cells and Caki2 cells expressing inducible PBRM1 were plated on day 0 and cells were harvested on designated time points for analysis. A designation of **** = p<0.0001 (paired Student *t*-test). Error bars represent s.d. n = 4.
